# How smart was *T. rex*? Testing claims of exceptional cognition in dinosaurs and the application of neuron count estimates in palaeontological research

**DOI:** 10.1101/2024.01.10.575006

**Authors:** Kai R. Caspar, Cristián Gutiérrez-Ibáñez, Ornella C. Bertrand, Thomas Carr, Jennifer A. D. Colbourne, Arthur Erb, Hady George, Thomas R. Holtz, Darren Naish, Douglas R. Wylie, Grant R. Hurlburt

**Affiliations:** Institute of Cell Biology, Heinrich Heine University Düsseldorf, Düsseldorf, Germany; Department of Game Management and Wildlife Biology, Faculty of Forestry and Wood Sciences, Czech University of Life Sciences, Prague, Czech Republic; Department of Biological Sciences, University of Alberta, Edmonton, Canada; Institut Català de Paleontologia Miquel Crusafont, Universitat Autònoma de Barcelona, Barcelona, Spain; Department of Biology, Carthage College, Kenosha, Wisconsin, USA; Comparative Cognition Unit, Messerli Research Institute, University of Veterinary Medicine Vienna, Vienna, Austria; School of GeoSciences, Grant Institute, University of Edinburgh, Edinburgh, Scotland, UK; Center for Science, Teaching, and Learning, Rockville Centre, New York, USA; School of Earth Sciences, University of Bristol, Bristol, UK; Department of Geology, University of Maryland, College Park, Maryland, USA; Department of Paleobiology, National Museum of Natural History, Washington, DC, USA; School of Biological Sciences, Faculty of Environment and Life Sciences, University of Southampton, Southampton, United Kingdom; Department of Natural History, Royal Ontario Museum, Toronto, Ontario, Canada

**Author notes:** These authors contributed equally (Kai R. Caspar, Cristián Gutiérrez-Ibáñez, Grant R. Hurlburt).

**Keywords:** endocast, palaeoneurology, brain evolution, comparative cognition, graphic double integration

## Abstract

Recent years have seen increasing scientific interest in whether neuron counts can act as correlates of diverse biological phenomena. Lately, Herculano-Houzel (2023) argued that fossil endocasts and comparative neurological data from extant sauropsids allow to reconstruct telencephalic neuron counts in Mesozoic dinosaurs and pterosaurs, which might act as proxies for behaviors and life history traits in these animals. According to this analysis, large theropods such as *Tyrannosaurus rex* were long-lived, exceptionally intelligent animals equipped with “macaque- or baboon-like cognition” whereas sauropods as well as most ornithischian dinosaurs would have displayed significantly smaller brains and an ectothermic physiology. Besides challenging established views on Mesozoic dinosaur biology, these claims raise questions on whether neuron count estimates could benefit research on fossil animals in general. Here, we address these findings by revisiting Herculano-Houzel’s (2023) work, identifying several crucial shortcomings regarding analysis and interpretation. We present revised estimates of encephalization and telencephalic neuron counts in dinosaurs, which we derive from phylogenetically informed modeling and an amended dataset of endocranial measurements. For large-bodied theropods in particular, we recover significantly lower neuron counts than previously proposed. Furthermore, we review the suitability of neurological variables such as neuron numbers and relative brain size to predict cognitive complexity, metabolic rate and life history traits in dinosaurs, coming to the conclusion that they are flawed proxies of these biological phenomena. Instead of relying on such neurological estimates when reconstructing Mesozoic dinosaur biology, we argue that integrative studies are needed to approach this complex subject.

## Introduction

The Late Cretaceous North American theropod dinosaur *Tyrannosaurus rex* is a superlative predator, being among the largest, heaviest, and most powerful (in terms of bite force) terrestrial carnivores of all time (Gignac and Erickson 2017; Sakamoto 2022; Henderson, 2023). Recently, Herculano-Houzel (2023) proposed that anthropoid primate-level intelligence should be added to *T. rex*’s already impressive predatory resume based on high estimated numbers of telencephalic neuron counts in large-bodied theropod taxa. This conclusion emerged from a paradigm whereby neurological variables estimated from endocasts can, so it is claimed, be used to infer metabolic parameters, social behaviors, and longevity in fossil species. Here, we test whether this approach and its remarkable prospects withstand scrutiny.

The hypothesis of exceptional intelligence in dinosaurs such as *T. rex* challenges the consensus of crocodile-like cognition in these animals, a position informed by comparative anatomical data (Rogers, 1998; Witmer & Ridgely, 2009; Hurlburt et al., 2013). Moreover, this claim bears ramifications that extend beyond specialized biological disciplines due to its potential to create long-lasting impacts on the public’s perspective on dinosaurs, evolution, and the scientific process. Given the extreme contrast between Herculano-Houzel’s (2023) proposal and more traditional perspectives on dinosaur biology, we revisit the claim of exceptional intelligence in these animals through an assessment of her methodology and a reanalysis of the underlying data. By integrating perspectives from both paleontology and neontology, we evaluate the potential benefits and limitations of neuron count estimation in research on the behavior and physiology of fossil species. We begin with a brief review of dinosaur paleoneurology and a discussion of how Herculano-Houzel’s (2023) approach aims to expand the field’s methodological tool kit.

### Dinosaur paleoneurology and the prospects of neuron count estimates for the field

Paleoneurology is a subfield of paleontology dedicated to research on the nervous systems of extinct animals. Because soft tissues are not readily preserved in the fossil record, paleobiologists rely on endocasts when studying the brains of extinct species (Paulina-Carabajal et al. 2023). An endocast can be a natural (infilling), artificial (mold) or virtual (digitally reconstructed) cast of the endocranial cavity that is formed by the bones of the braincase.

The study of extinct species’ endocasts, including those of dinosaurs, can be traced back to the 1800s (e.g., Cuvier, 1812; Marsh, 1879). However, the field was truly defined by Edinger (1929) who effectively introduced the concept of geological time to neurobiological studies. Before her, anatomists made comparisons between endocasts and fresh brains, but without considering the stratigraphic context (Buchholtz & Seyfarth, 2001). Jerison (1973) built on Edinger’s work by studying brain evolution in a quantitative manner and developed the encephalization quotient (EQ) as an estimate of relative brain size, applicable to both extant and extinct species. Later, the advent of X-ray computed tomography at the end of the 1990’s transformed the field and provided novel ways in which the neurosensory systems of extinct species could be studied (e.g., Knoll et al., 1999; Witmer et al., 2008). Despite these crucial innovations, however, paleoneurology has so far remained largely restricted to the measurement and comparison of gross morphology, limiting our understanding of how the brains of Mesozoic dinosaurs and other extinct animals worked.

Pterosaurs and dinosaurs (the latter including birds) form the clade Ornithodira (Fig. 1), the closest extant relatives of which are crocodilians (Fig. 1). Together, both lineages, which separated about 250 million years ago, comprise the clade Archosauria (e.g., Legendre et al., 2016). Next to birds, crocodilians therefore represent a critical reference point in reconstructing the nervous systems of extinct ornithodirans.

**Figure 1:**
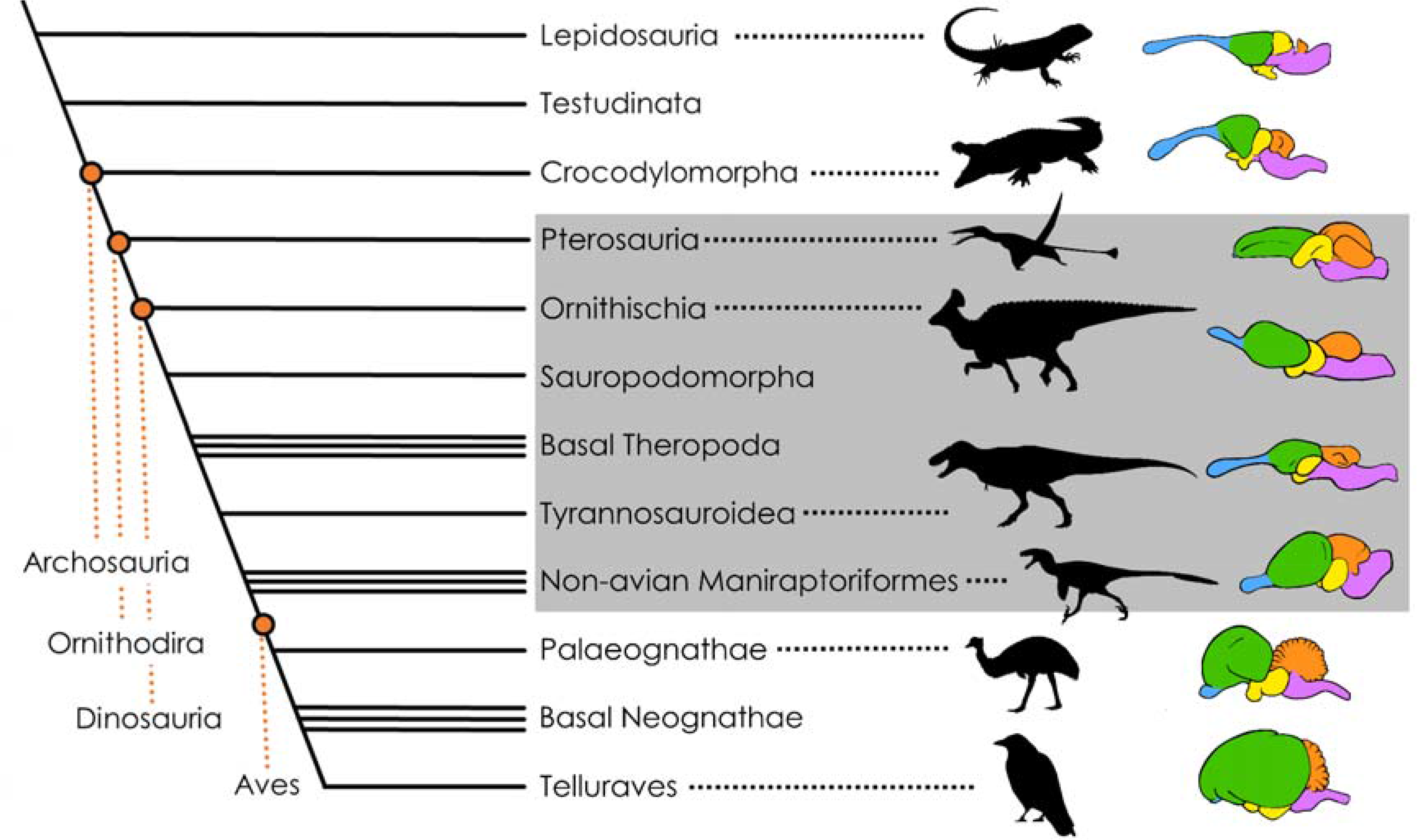
Simplified phylogeny of the Sauropsida (= total group Reptilia) with a focus on the taxon Ornithodira (the least inclusive clade containing pterosaurs and dinosaurs, see revised definition of Nesbitt, 2011) and representative color-coded brain morphologies, excluding the pituitary (not to scale). Blue: olfactory bulb and tracts, Green: pallium (homologous to the mammalian cerebral cortex), Orange: cerebellum, Yellow: diencephalon and optic tectum; Violet: brain stem. Olfactory structures, pallium and subpallium comprise the telencephalon. The gray overlay indicates extinct taxa, the brain morphologies of which are approximated. Note that the brain morphology in *T. rex* and its relatives (Tyrannosauroidea) is conspicuously plesiomorphic when compared to the other ornithodirans pictured here (see e.g., Giffin, 1989). Definitions of notable dinosaur clades: Ornithischia - a large group of primarily herbivorous dinosaurs, excluding the long-necked sauropodomorph dinosaurs, defined as the most inclusive clade including *Triceratops* but not *Diplodocus* nor *Tyrannosaurus*. Most popular representatives of this group include horned or otherwise heavily armored forms such as *Triceratops*, *Stegosaurus* and *Ankylosaurus* as well as the hadrosaurs, colloquially known as duck-billed dinosaurs. Sauropodomorpha - the long-necked and often particularly large-bodied herbivorous dinosaurs, defined as the most inclusive clade including *Diplodocus* but not *Triceratops* nor *Tyrannosaurus;* Theropoda - the bipedal, mostly carnivorous dinosaurs, the most inclusive clade including *Tyrannosaurus* but not *Diplodocus* or *Triceratops*. The birds are part of this clade (see Baron et al., 2017 for definitions of Ornithischia, Sauropodomorpha and Theropoda); Tyrannosauroidea, the most inclusive clade of theropods containing *Tyrannosaurus* but not more bird-like taxa such as *Velociraptor* and *Ornithomimus* (Sereno *et al*., 2009); Maniraptoriformes, the least inclusive clade containing *Velociraptor* and *Ornithomimus* but not earlier-diverging theropods like *Tyrannosaurus* (Holtz, 1996). Silhouettes were taken from PhyloPic (listed from top to bottom): *Morunasaurus* (in public domain), *Crocodylus* (in public domain), *Rhamphorhynchus* (by Scott Hartman), *Olorotitan* (by ДиБгд, vectorized by T. Michael Keesey), *Tyrannosaurus* (by Matt Dempsey), *Dromaeosaurus* (by Pranav Iyer), *Dromaius* (by Darren Naish), *Corvus* (in public domain).

Interestingly, highly disparate patterns of endocranial tissue organization are realized in these two extant clades. One fundamental difference relates to the portion of the endocranial cavity which is occupied by the brain rather than by the associated meningeal tissues (including the dura mater and arachnoid mater) and cerebrospinal fluid (Fig. 2). In crocodilians, nervous tissue only fills a fraction of the braincase (Hopson, 1979; Jirak & Janacek, 2017; Watanabe et al. 2019). Longitudinal venous sinuses course along the dorsal and ventral aspect of the brain, obscuring its true shape in casts of the braincase.

**Figure 2:**
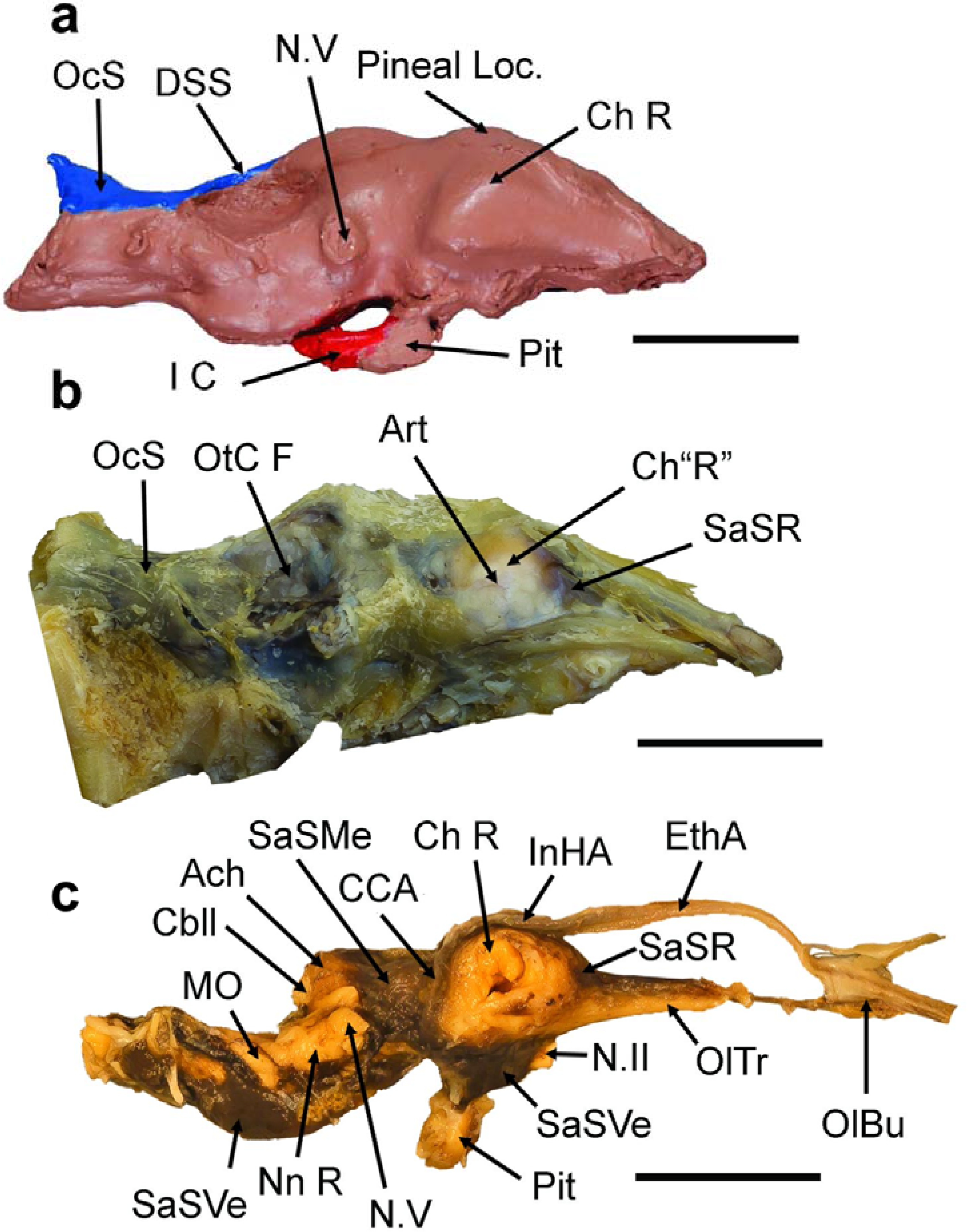
The endocast and endocranial tissue organization in the American alligator (*Alligator mississippiensis*), illustrating the plesiomorphic condition within the clade Archosauria. Scale bar = 2 cm in all cases. **a:** Endocast of a wild *A. mississippiensis* (Fla.F&G.HarvestTag 937095), Dorsal cranial length (DCL), 342.90 mm, right lateral view. Reduced in size to match proportions of brain in **b** & **c**. **b**: Dura mater around the brain of *A. mississippiensis*, specimen CITES FLM 12-29409, DCL, 380 mm, left lateral view (reversed). **c**: Brain within arachnoid of FLM 12-29409. Brown-red material is dried blood filling the subarachnoid space (**SaS**), right lateral view. **Abbreviations: Ach**, arachnoid mater (covering the cerebellum); **Art**, artery on external wall of dura mater over the lateral pole of the cerebrum; **Cbll**, cerebellum; **CCA**, caudal cerebral artery; **Ch L** or **R**, left or right cerebral hemisphere; **DSS**, dorsal sagittal sinus; **EthA**, common ethmoid artery; **I C**, internal carotid artery; **InHA**, interhemispheric artery; **MO**, medulla oblongata; **N.II**, optic nerve; **N.V**, (cast of) trigeminal nerve; **Nn R**, roots of nerves IX-XI; **OC**, occipital condyle; **OcS**, occipital sinus; **OlBu** & **Tr**, olfactory bulb & tract; **OtC F**, fossa of otic capsule; **Pineal Loc**, pineal gland location; **Pit**, pituitary gland; **SaSMe**, mesencephalic subarachnoid space; **SaSR**, rostral SaS; **SaSVe**, ventral SaS; **SN.I**, first spinal nerve. The rostral end of the cerebrum is below the arrow for **SaSR** in **B**. Both specimens are housed in the private collection of G. R. Hurlburt.

Furthermore, the size of the brain relative to both the endocranial volume and total body size, decreases during crocodilian ontogeny, even over the course of adulthood (Hurlburt et al., 2013; but note that absolute brain volume increases with body size, even in adults - Ngwenya et al., 2013).

Endocast morphology indicates that the endocranial cavity in most non-avian dinosaurs was organized in crocodilian-like fashion and comparative studies suggest that this configuration was indeed ancestral for the clade Archosauria (Witmer et al., 2008; Hurlburt et al., 2013; Fabbri & Bhullar, 2022). For tyrannosauroids specifically, which are among the best-studied dinosaurs when it comes to palaeoneurology, endocasts representing different ontogenetic stages suggest that brain size (relative to endocranial volume) decreased with age (Brusatte et al., 2009; Witmer & Ridgely, 2009; Bever et al., 2013), as is the case in modern crocodilians. As in, crocodilians, most dinosaurian endocasts do not faithfully capture the volume and anatomy of the brain, particularly its posterior regions such as the cerebellum (Watanabe et al., 2019). This contrasts with the situation in birds and most mammals for which endocasts represent excellent brain size proxies (e.g., Iwaniuk and Nelson, 2002; Bertrand et al., 2022).

The avian pattern probably evolved at the root of the theropod dinosaur clade Maniraptoriformes, which includes ornithomimosaurs (‘ostrich-mimic’ dinosaurs) and maniraptorans (the bird-like oviraptorosaurs, dromaeosaurids and kin, and birds themselves) (Osmólska, 2004; Balanoff et al., 2013; Fig. 1). Maniraptoriform brains have enlarged cerebral and cerebellar regions that almost fully occupy the endocranial cavity, as evidenced by brain contours faithfully captured by the endocranium and extensive vascular imprints. There is no evidence that the brains of other dinosaurs similarly contacted the endocranial surface (pachycephalosaurs pose an exception to this pattern but are not covered in this article, their endocranial anatomy is described in Hopson, 1979, Giffin, 1989, and Evans, 2005; we discuss other suggested cases of secondarily increased endocranial fills in dinosaurs in Supplementary File 1). Pterosaurs are similar to maniraptoriforms in also possessing brains that fit tightly into the endocranial cavity (Witmer et al., 2003).

Aside from general endocranial tissue organization, the neuroarchitecture and circuitry of the forebrain in birds and crocodilians differs notably from one another (Ulinski & Margoliash, 1990; Briscoe et al., 2018; Briscoe & Ragsdale, 2018). Comparisons with other sauropsids demonstrate that again the crocodilian condition is more plesiomorphic (Briscoe & Ragsdale, 2018). To which extent non-avian dinosaurs and pterosaurs resembled the two extant archosaur groups in these regards cannot be reliably reconstructed, since they lack osteological correlates.

The inferred brain anatomy of various dinosaur groups has been discussed elsewhere (Paulina-Carabajal et al. 2023) and reviewing it here is beyond the scope of this article. We aim instead to focus on what endocast-based methods potentially reveal about the behavior and cognition of extinct species. While considering the aforementioned limitations, endocasts from fossil ornithodirans allow us to reasonably estimate basic neuroanatomical measures such as EQ, as well as to deduce specific sensory specializations (e.g., Witmer et al., 2003; Witmer & Ridgely, 2009; Zelenitsky et al., 2011). Nonetheless, it is generally assumed that the predictive power of these data in elucidating the cognitive capacities of a fossil species is low (Paulina-Carabajal et al., 2023). Researchers have long sought to identify robust morphological correlates of cognition but have found traditional proxies such as EQ and absolute brain size to be limited and problematic regarding their conceptual justifications (Van Schaik et al., 2021). Current debates focus on whether refined neuroanatomical measures such as “cognitive brain size” (Van Schaik et al., 2021) and brain region-specific neuron counts (Herculano-Houzel, 2011; Kabadayi et al., 2016; Logan et al., 2018; Sol et al., 2022) might be able to overcome these issues. The quantification of these, however, seemed out of reach for vertebrate palaeontology.

With this in mind, the approach proposed by Herculano-Houzel (2023) is of great potential significance: It entails that endocasts of extinct taxa can be used to model neuron counts if neurological data from related extant species can be taken into account. If valid, this technique would potentially allow researchers to elucidate aspects of brain physiology that cannot be inferred from endocast morphology alone. Herculano-Houzel & Kaas (2011) and Herculano-Houzel et al. (2011) pioneered this approach for fossil hominins and extinct giant rodents, but Herculano-Houzel (2023) was first in applying this methodology to fossil sauropsid groups separated from their extant relatives by hundreds of millions of years of evolution, namely pterosaurs and Mesozoic dinosaurs.

Indeed, Herculano-Houzel (2011, 2017, 2023) has argued emphatically that neuron counts represent reliable estimates for cognitive abilities in extant vertebrates, markedly outperforming other measures such as relative or absolute brain size. If we accept this premise, accurate modeling of neuron counts in dinosaurs based on endocast volumes and comparative neurological data might appear as a promising new method to elucidate the behavior and cognitive capacities of these animals.

### The methodology and rationale of Herculano-Houzel (2023)

Herculano-Houzel (2023) reconstructed relative brain size and neuron counts for 29 dinosaur and pterosaur species based on comparative data from extant non-avian and avian sauropsids (“reptiles” and birds respectively; Olkowicz et al., 2016; Kverková et al., 2022). Although we want to avoid lengthy discussions about taxonomy, it is worth noting that some of these are no longer considered valid taxonomic entities (see below; an updated nomenclature for relevant dinosaurs is included in Tab. 1). For instance, *Rhamphorhynchus muensteri* and *R. gemmingi* have long been synonymized (Bennett, 1995). Surprisingly, Herculano-Houzel (2023) inferred an ectothermic metabolism for one, and endothermy for the other based on assumptions about relative brain size.

**Table 1:**
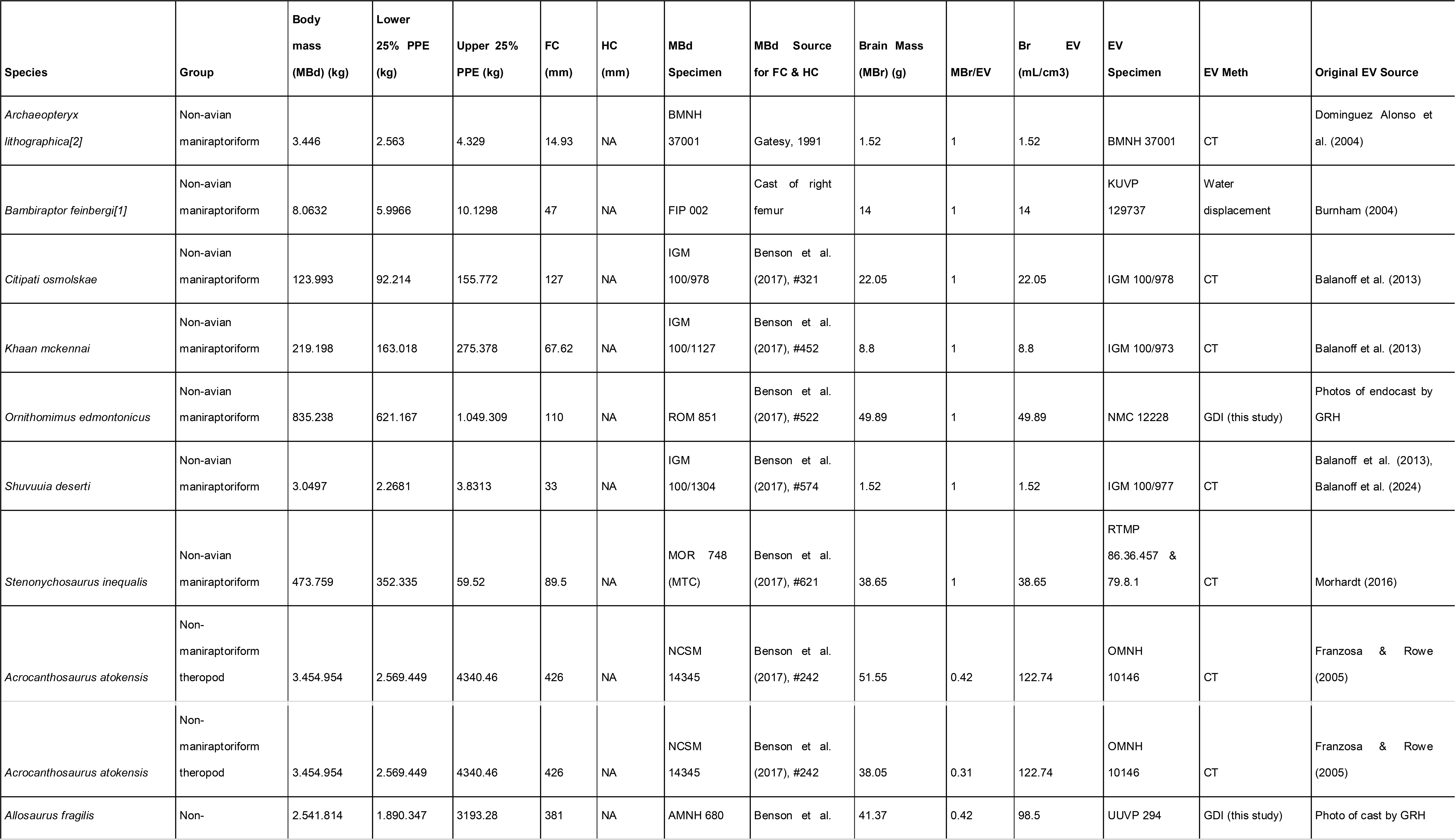
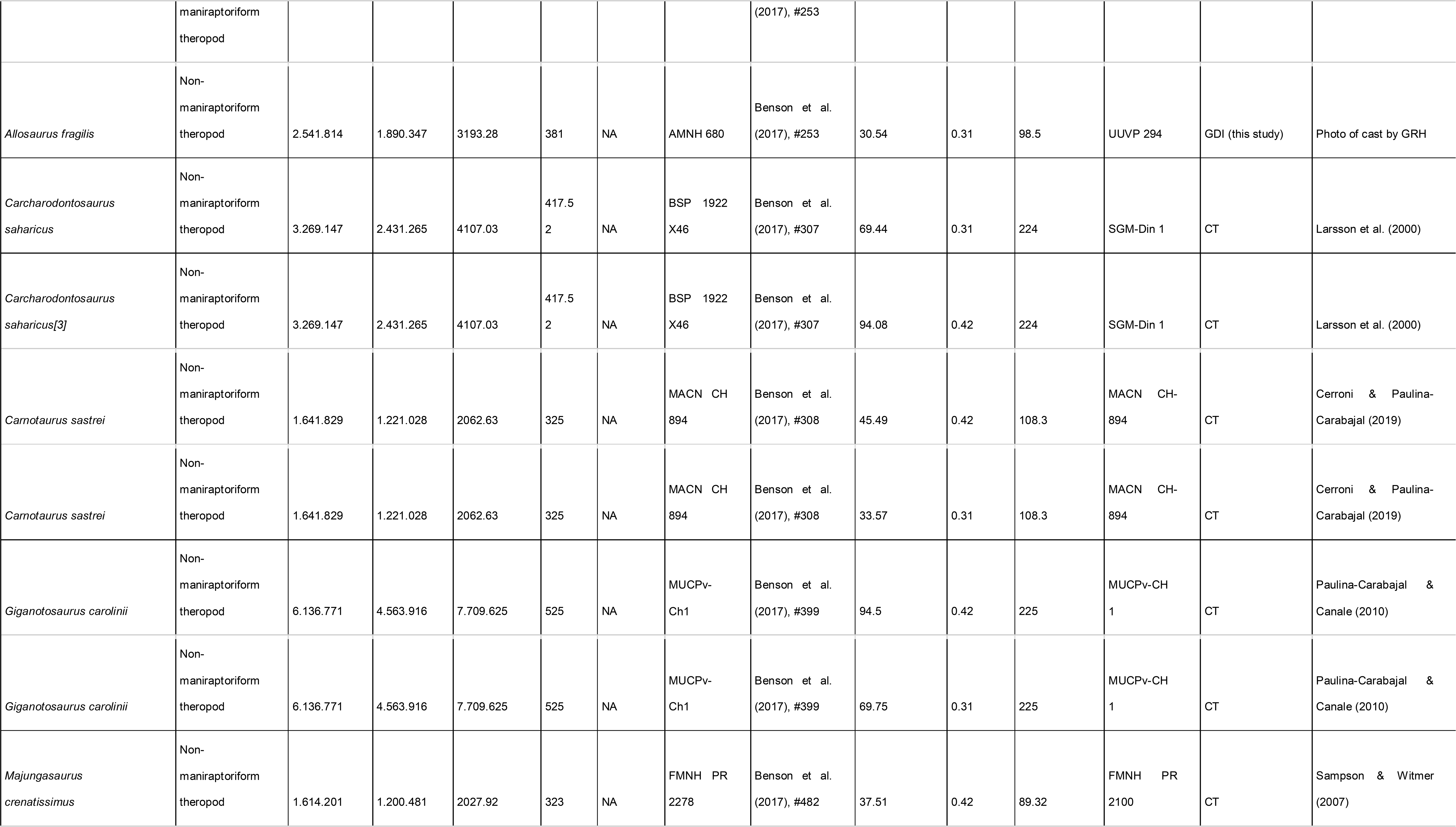
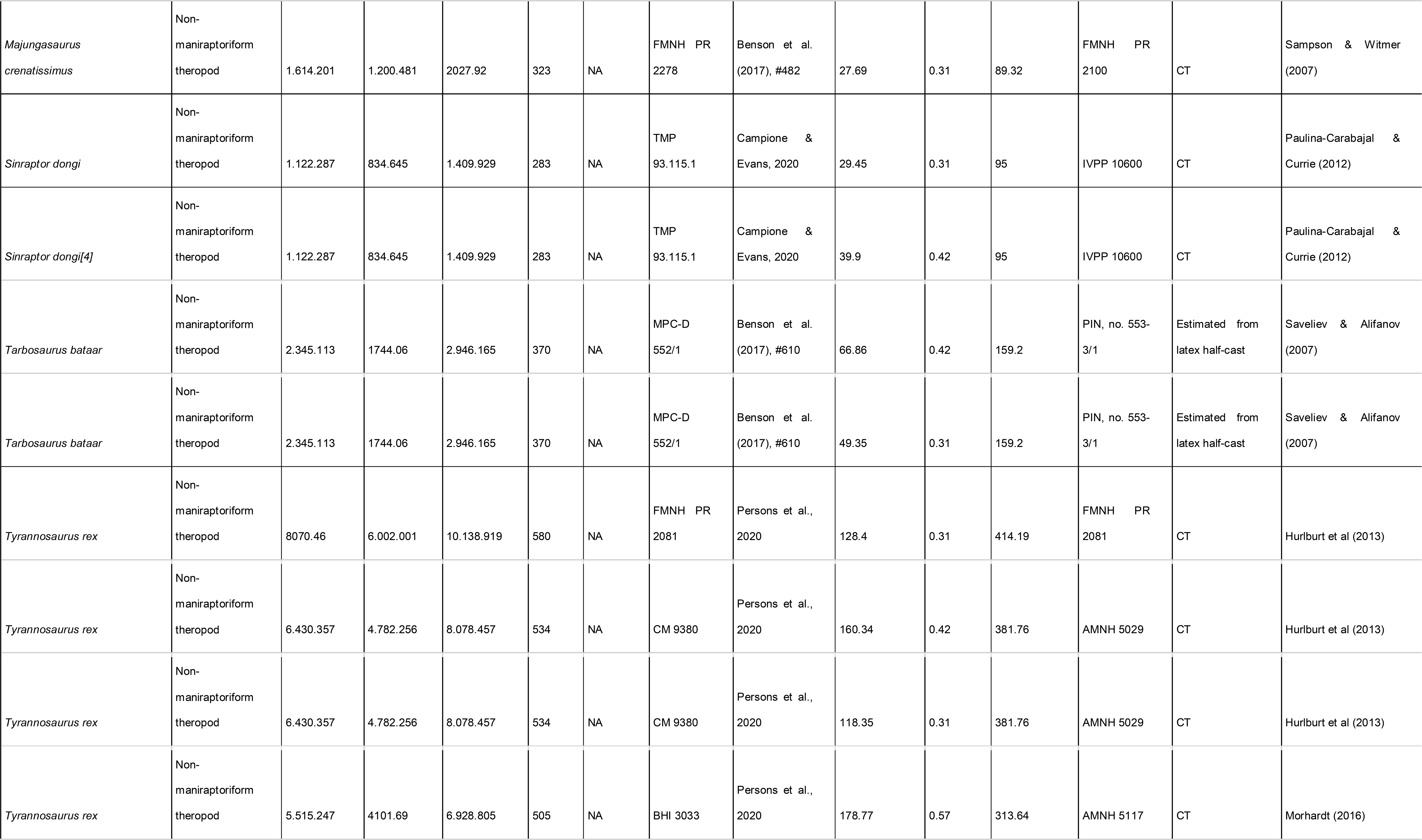
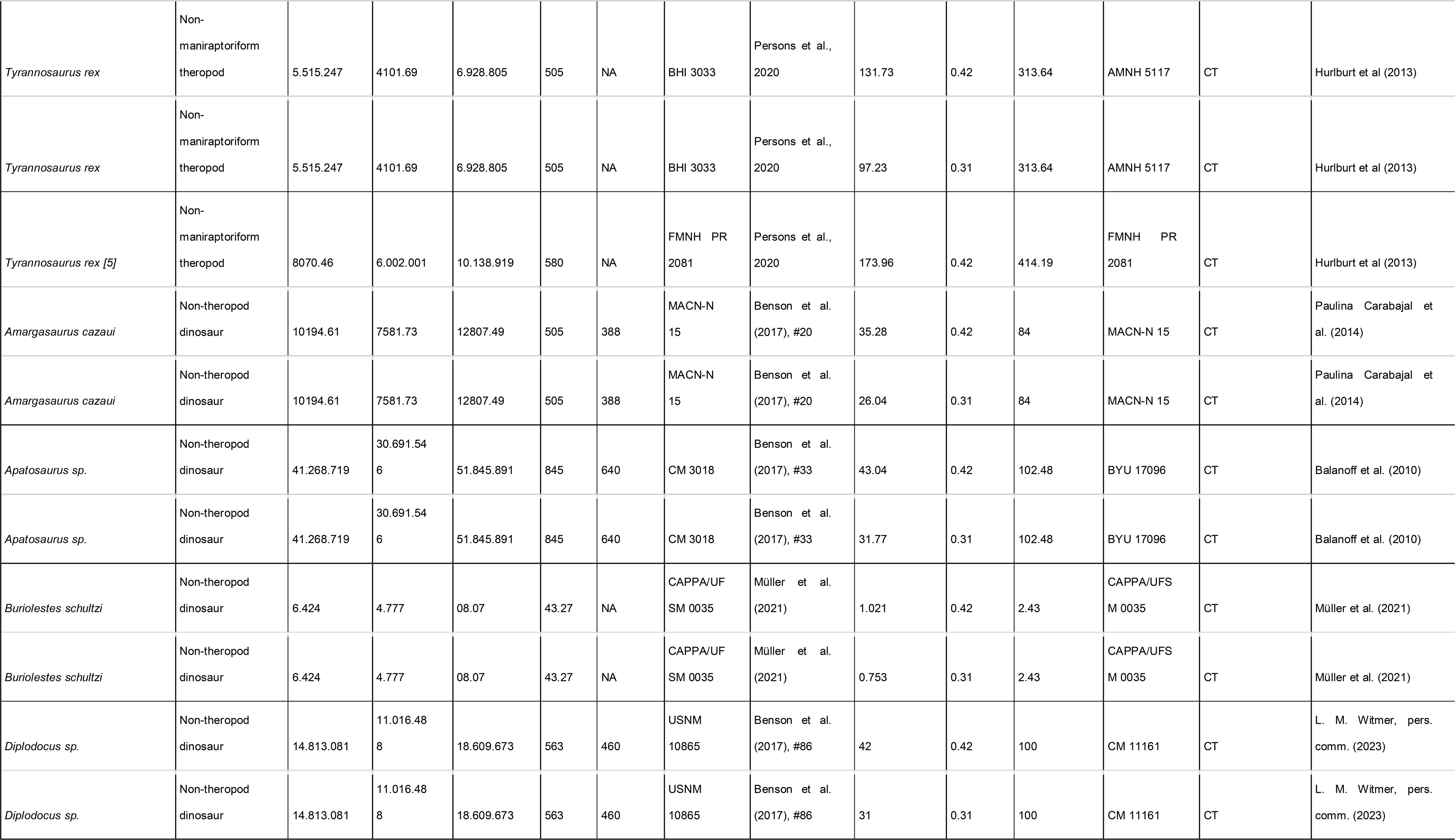
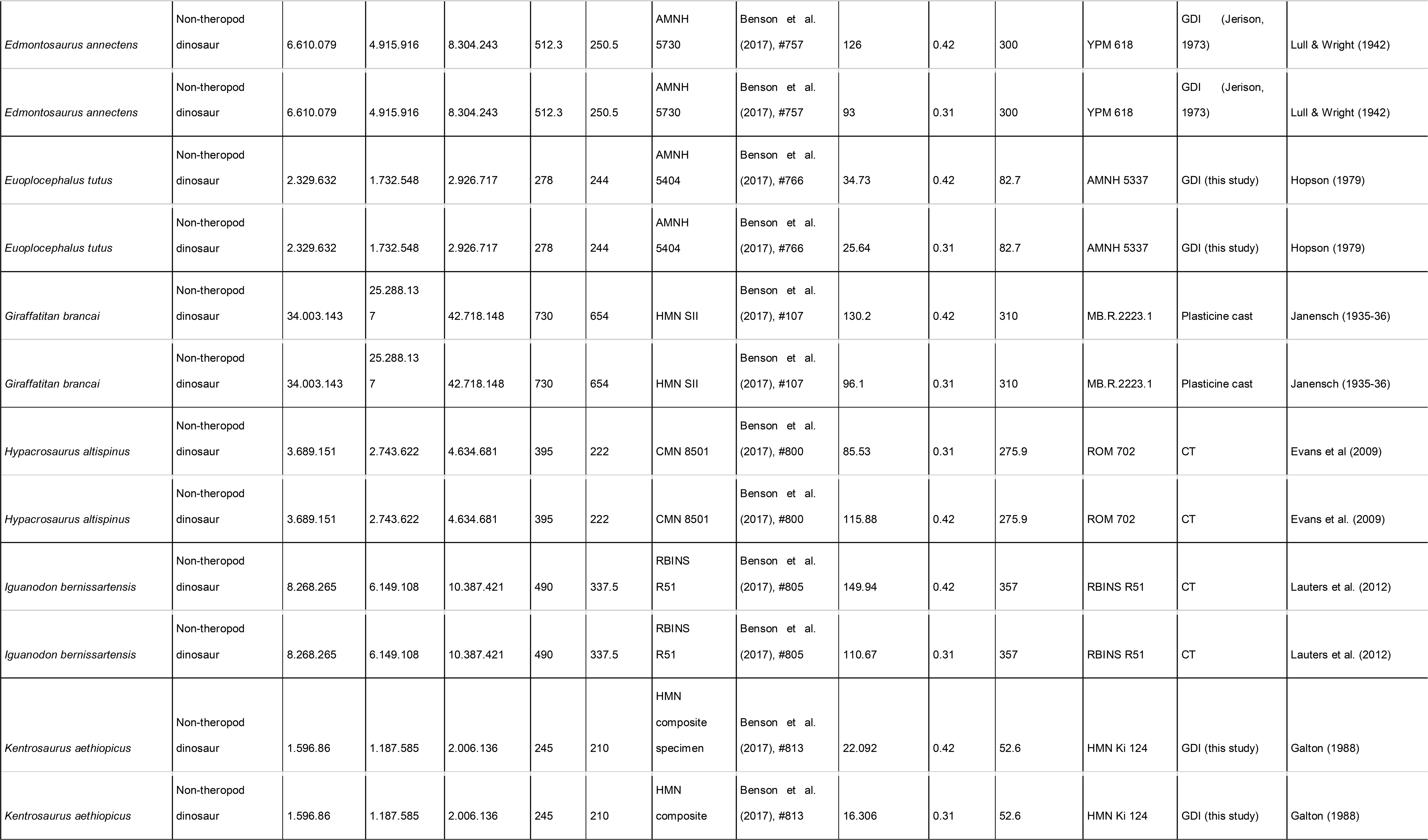
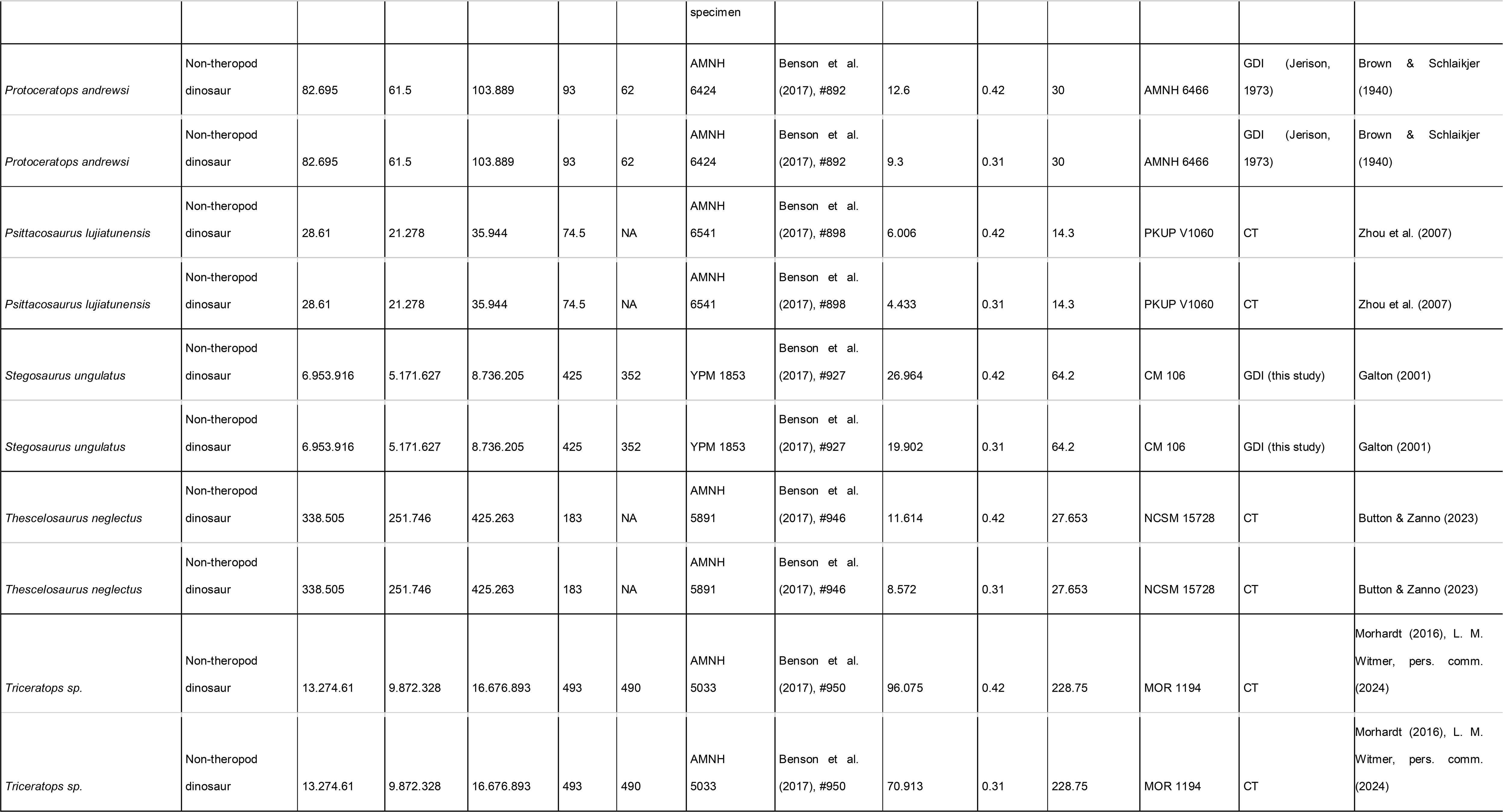
Estimates of brain (MBr, g; derived from endocranial volume, EV) and body mass (MBd, kg) in Mesozoic dinosaurs. For maniraptoriform theropods, we assumed that brain endocast volume equals brain volume. For other dinosaurs, we assumed a brain:endocast ratio of 31 - 42 %. Body mass was calculated based on stylopodial circumference (femoral circumference (FC) for bipedal and femoral as well as humeral (HC) circumference for quadrupedal species). **Footnotes: [1]** *Bambiraptor feinbergi*. MBd from femur circumference, measured on a cast of the right femur of an adult specimen now in the collection of the Vertebrate Paleontology Division, ROM (FIP 002). Original elements of this specimen are now in the collection of the AMNH. **[2]** *Archaeopteryx lithographica*. MBd from FC (14.93 mm) estimated from femur length (60.5mm) of BMNH 37001 (Gatesy, 1991). FC was calculated using the equation from Benson et al. (2017): log10(Femur circumference estimate from femur length) = 1.132 * log10(Femur length) - 0.8429. **[3]** *Carcharodontosaurus saharicus.* MBd from estimated femur circumference (FC = 417.52 mm) from Benson et al. (2017) for BSP 1922 X46. The specimen has been destroyed and is no longer accessible. **[4]** *Sinraptor dongi*. MBd based on FC (283 mm) from Campione & Evans (2020), measurement taken from TMP 93.115.1 (cast of IVPP 10600). Benson et al. (2017) consider IVPP 10600 a subadult based on reported incomplete fusion of cervical vertebrae; However, Paulina-Carabajal & Currie (2012) noted that the degree of cranial suture fusion indicates that the specimen is an adult or at least a large subadult. We consider it an adult here. **[5]** *Tyrannosaurus rex* specimens. MBd based on FC values listed by Persons et al. (2020). For FMNH PR 2081 (“Sue”), but not the other individuals considered here, both FC and EV are available. We associated the EV (313.636 mL) of AMNH 5117 with the MBd (5515 kg) of BHI 3033 (“Stan”), as both specimens have been considered proxies for each other (G. M. Erickson, pers comm. to G.R. Hurlburt, 2005). The EV (381.8 mL) of AMNH 5029 is here linked to the MBd (6430 kg) of CM 9380 (holotype specimen) because it fell between MBd’s associated with EVs of FMNH PR 3081 and AMNH 5117. Besides our brain:endocast ratios, we also apply the 57% ratio proposed by Morhardt (2016) to this species.

Neuron count estimates for fossil taxa only concerned the telencephalon, a major brain region which is critically involved in cognitive and motor functions as well as the processing of sensory information. It encompasses the pallium (which is homologous to the cerebral cortex in humans and other mammals) and subpallium as well as the olfactory bulbs and tracts (Fig. 1). To understand the rationale behind the approach of reconstructing neuron counts in fossil species, two important matters must be pointed out: firstly, among jawed vertebrates, body size and brain size are highly correlated, exhibiting a constant allometric relationship overall (Tsuboi et al., 2018). It should be noted however, that scaling relationships can vary to some extent between major taxa as well as between early- and late-diverging members of a clade (Ksepka et al., 2020; Bertrand et al., 2022). Secondly, neuronal densities (the number of neurons in a given volume of nervous tissue) can differ profoundly between different vertebrate taxa. Based on current evidence, the highest neuron densities among land vertebrates are found in the bird clade Telluraves, consisting of birds of prey, rollers, parrots, songbirds and kin, while the lowest occur among crocodilians and turtles (Kverková et al., 2022). For instance, the goldcrest (*Regulus regulus*), short-tailed shrew (*Blarina* sp.) and painted turtle (*Chrysemys picta*) have brains of equal mass (ca. 0.37 g), but there is remarkable disparity in their whole brain neuron numbers, which range from 14.3 M in the turtle over 58.8 M in the shrew to 164 M in the passerine bird (Olkowicz et al., 2016; Kverková et al., 2022). This example illustrates that brain size alone is not a reliable predictor of neuron counts across distantly related clades (compare Herculano-Houzel et al., 2014; Olkowicz et al., 2016), which makes their inference in fossil groups inherently difficult.

To decide which neuronal density patterns apply to specific groups of dinosaurs and pterosaurs, Herculano-Houzel (2023) relied on brain x body mass regressions. The brain and body mass datasets used were taken from various literature sources and, as we attempt to show here, both are problematic. In the resulting regression plot, she identified theropods clustering distinctly from most other species. On average, they appeared to exhibit larger brains for a given body size than the remaining dinosaur or pterosaur taxa. When comparing the regression lines for extinct groups with those of living birds on the one hand and squamates and turtles on the other, Herculano-Houzel (2023) noted that the theropod regression fits with the avian one, while the remaining ornithodiran groups aligned more with the non-avian sauropsid regression line.

Based on these analyses, the author made two critical assumptions: First, since only theropod brain-body data aligned with those of endothermic extant sauropsids, namely birds, the other groups (aside from specific pterosaurs and ornithischians that cluster with theropods) should be considered ectothermic. Second, telencephalic neuron densities in theropod brain tissue should have been comparable to those found in certain extant bird taxa (that is, to those found in a polyphyletic assemblage denoted as “pre-K-Pg birds” that includes Palaeognathae, Galloanserae and Columbiformes and which is considered to form a neurological grade: Kverková et al., 2022), whereas those of the other groups should resemble densities encountered in squamates and turtles. No further justification for these suggestions is provided.

Applying the avian scaling regime, Herculano-Houzel (2023) estimated remarkably high telencephalic neuron counts in large-bodied theropods such as *Acrocanthosaurus atokensis* (2.1 billion) and *T. rex* (3.3 billion) which would exceed those of any extant bird and be comparable to large-bodied Old World monkeys such as baboons (*Papio anubis* - Olkowicz et al., 2016). Based on this apparent similarity to anthropoid primates, she further speculated that these giant theropods would have crafted and used tools and exhibited cultural behaviors (Herculano-Houzel, 2023).

We regard the methodology of Herculano-Houzel (2023) as problematic and disagree with her physiological and behavioral interpretations. Before we attempt to replicate her findings with a more refined analytical approach, we want to enumerate the most important flaws of the article and how they affect the inferences made.

### Issues with Herculano-Houzel’s method and rationale

A key problem for paleoneurology lies in the fact that an endocast does not necessarily reflect the morphology of an animal’s brain. As discussed in previous sections, the endocasts of most non-avian dinosaurs differ markedly in size and shape from the actual brain, as is the case in crocodilians (Fig. 1). Unfortunately, not all studies from which Herculano-Houzel (2023) derived her raw data considered this issue (see below). In addition, the percentage of endocranial space filled by the brain, as well as its proportions may be further influenced by ontogeny (Bever et al., 2013; Hurlburt et al., 2013; Ngwenya et al., 2013; Jirak & Janacek, 2017; Hu et al., 2021). The latter point is relevant because Herculano-Houzel (2023) included several specimens which corresponded to juveniles rather than adults, and thus might have introduced biases to the dataset (see below). Interestingly, at least in crocodilians, neuronal densities in the brain are also affected by ontogenetic stage (Ngwenya et al., 2016).

To arrive at the estimated telencephalic neuron count of > 3 billion for *T. rex*, Herculano-Houzel (2023) assumed a brain mass of 343 g. However, this presupposed that endocast volume equaled brain volume in this species. While it has indeed been claimed that in theropods such as *T. rex*, the brain filled the entire endocranial cavity (Balanoff et al., 2013), this hypothesis is, as previously discussed, contradicted by multiple lines of evidence. More conservative inferences suggest a brain mass of approximately 200 g (Hurlburt, 1996; Hurlburt, et al, 2013; Morhardt, 2016) or possibly even lower (this study; Tab. 1) for *T. rex*. Herculano-Houzel (2023) acknowledged these lower estimates but chose to rely on the inflated values for large theropod brain masses in accompanying figures and the article’s Discussion section.

Moreover, while the literature-derived brain mass estimates used for their analyses did in some cases include the olfactory tracts and bulbs (Franzosa & Rowe, 2005, Balanoff et al., 2013), these were omitted in others (Hurlburt 1996; Hurlburt et al., 2013). This incongruence created critical biases, affecting both the inference of telencephalic neuron counts and relative brain size estimates. The latter are additionally skewed by the fact that body masses used by Herculano-Houzel (2023) were not determined via a uniform methodology but compiled from sources applying various approaches. There are several ways to estimate body mass in extinct animals and they can differ greatly regarding their outcomes and precision (Campione & Evans, 2020). When compared to body mass estimates derived from stylopodial circumference, a well-established and robust method (Campione & Evans, 2020), some striking differences become apparent (Tab. 1; Herculano-Houzel, 2023).

Another flaw of Herculano-Houzel’s (2023) approach is the neglect of brain morphology to inform its analyses. To estimate telencephalic neuron numbers in fossil species, the mass of the telencephalon needs to be approximated first. For theropods, Herculano-Houzel (2023:6) extrapolated this variable from extant bird data while stating that “within a clade, brain mass has strongly predictive power to arrive at estimates of numbers of telencephalic neurons in a brain of known mass, once the neuronal scaling rules that presumably apply are known.” However, this statement can only hold true if the general proportions of the telencephalon compared to the remaining brain are roughly constant in the group of concern, which is a precondition that Herculano-Houzel (2023) did not test for in the fossil sample. Indeed, avian brains only poorly reflect the brain morphologies found in the majority of Mesozoic dinosaurs (reviewed by Paulina-Carabajal et al. 2023) and their general proportions are only comparable to those found among maniraptoriform theropods (Balanoff et al., 2013; Fig. 1). An important difference concerns the pallium, which crucially contributes to higher cognitive functions, and greatly increased in size within the maniraptoriform radiation (Balanoff et al., 2013). The same is true for the cerebellum, a part of the brain which is not encompassed by the telencephalon but is also involved in various aspects of cognition in amniotes (Spence et al., 2009). Thus, the telencephalic mass and proportions of non-maniraptoriform theropods, such as *T. rex*, cannot be adequately modeled based on extant birds. Similar limitations need to be considered when reconstructing traits of, for instance, the pterosaur or sauropodomorph telencephalon based on extant non-avian sauropsids and they also apply to our own empirical approach.

Herculano-Houzel (2023) hypothesized that the inferred incongruence in relative brain size between theropods and other dinosaurs reflects differences in thermobiology, which would justify applying avian neuronal scaling schemes to the former and non-avian sauropsid scaling to the latter. Sauropodomorphs as well as selected ornithischians and pterosaurs are instead suggested to be ectothermic due to their relatively smaller brains. Both of these assumptions are problematic: First, multiple lines of evidence suggest that ornithodiran endothermy evolved long before theropods emerged and was likely already present in the Early Triassic common ancestor of dinosaurs and pterosaurs (e.g., Benton, 2012; Grigg et al., 2022). We will revisit this evidence and how it challenges the brain size-derived hypothesis in more detail in the Discussion section of this paper. Herculano-Houzel (2023) only referenced a single article on dinosaur thermobiology, that of Wiemann et al. (2022), to defend her standpoint on the matter. The study in question applies a promising but novel technique to infer endothermy based on lipoxidation end products in fossil bone that still has to prove itself. While it indeed suggests lowered metabolic rates in some ornithischians, it also infers an endothermic metabolism for pterosaurs and sauropodomorphs (Wiemann et al., 2022). Thus, its findings do not align with Herculano-Houzel’s (2023) assumptions.

Second comparisons between groups of extant vertebrates, especially birds and mammals, strongly suggest that there is no uniform relationship between neuron density and relative brain size or elevated metabolic rates (Kverkova et al., 2022; see also Estienne et al., 2024). We will elaborate on this aspect in the Discussion section but would like to state at this point already that it is not straight-forward to assume avian neuron densities in Mesozoic theropods simply because they exhibited endothermy or a potential increase in relative brain size. On the other hand, the extensive evidence for endothermy in other dinosaurs and pterosaurs does not entail that these groups could not have had neuron densities similar to those found in extant ectothermic sauropsids.

Other issues relate to the statistical methods employed by Herculano-Houzel (2023). Despite dealing with a large multi-species dataset, their analyses did not take phylogeny into account, which can produce mathematical artifacts. Phylogenetic relationships among taxa need to be statistically addressed because shared ancestry can result in non-independence of species-specific data points (Revell et al., 2008). Such non-independence is known as the phylogenetic signal, and it has been prominently recovered for relative brain size in extant sauropsids (Font et al., 2019). Hence, phylogenetically-informed modeling is necessary for adequately analyzing such datasets (Font et al., 2019).

In light of these substantial shortcomings, we attempt to replicate the findings of Herculano-Houzel (2023) with phylogenetically informed models of telencephalic neuron counts in fossil dinosaurs that acknowledge the issues lined out above. Different from her, we do not include pterosaurs into our analysis due to difficulties with estimating their body mass (especially for taxa with incomplete postcrania such as *Scaphognathus*) and because of the unclear taxonomic and ontogenetic status of some of the few available endocasts.

### Empirical part: modeling neurological variables in dinosaurs

#### Endocast sample composition, with notes on endocranial volumes provided by Hurlburt (1996)

We estimated the mass of the brain (MBr, g; excluding the olfactory tracts and bulbs) as well as its size relative to body mass (MBd, g) in 31 Mesozoic dinosaur taxa for which data on endocranial volume (EV, ml) have been published (Tab. 1; Suppl. File 1). Note that this study does not aim to provide a comprehensive dataset of dinosaur brain sizes. Given the questions we want to address, we focus on large-bodied theropods and taxa covered in previous comparative analyses. We included one endocast per species, except for *T. rex*, for which three adult endocasts (AMNH 5029, AMNH 5117, FMNH PR 2081) were considered. We only considered species for which we could calculate body mass based on stylopodial circumference (see below) to reliably infer encephalization. Due to this, our analysis does not cover all dinosaur species for which complete endocasts are available, nor all species that Herculano-Houzel (2023) included in her analyses (namely *Conchoraptor gracilis*, *Tsaagan mangas*, *Zanabazar junior*, and unnamed troodontid IGM 100/1126). Juvenile specimens considered by that study (*Alioramus altai* IGM 100/1844, *Gorgosaurus libratus* ROM 1247, and *Tyrannosaurus rex* CMN 7541 *= “Nanotyrannus lancensis”)* were omitted in this analysis to eliminate the confounding variable of ontogeny.

The only juvenile we include is *Bambiraptor feinbergi* KUVP 129737, which is one of the few maniraptoriform theropods that we can take into account. For this species, an adult femur (FIP 002) is available, allowing us to estimate body mass in fully grown individuals. Our method of body mass inference suggests that KUVP 129737 had attained about 45% of adult body mass when it died. Data from similar-sized extant rheas (*Rhea americana*), palaeognath birds which are close neuroanatomical analogs to highly derived theropods such as *Bambiraptor* (Balanoff et al., 2013), suggest that adult brain mass is already approached at that point of somatic development (Picasso et al., 2011; Picasso, 2012;). We therefore combine the juvenile endocranial measurement of *Bambiraptor* with adult body mass estimates.

Just as Herculano-Houzel (2023) did, we derive a significant portion of our EV values from Hurlburt (1996). However, many EV figures communicated in this reference must be considered outdated or otherwise flawed and were carefully bypassed here. We give detailed reasons for discarding or modifying data from Hurlburt (1996) and the references provided therein (Jerison, 1973; Hopson, 1979) in Suppl. File 1 Part A. Given that EVs from this problematic dataset are still widely used (e.g., Müller et al. 2021; Button & Zanno, 2023), we consider their revision an important aspect of this study. In cases where EVs appeared doubtful but appropriate illustrations or photographs of specimens were available, we recalculated EV using manual graphic double integration (GDI; see below for methodology). This was done for four species (*Allosaurus fragilis, Euoplocephalus tutus, Kentrosaurus aethiopicus & Ornithomimus edmontonicus*; see Suppl. File 1 for details on specimens).

### Brain mass estimates

We estimated the brain mass (MBr, g) of fossil dinosaurs from endocast volume (EV, mL). Because the specific gravity (density) of living amniote brain tissue approximates one (1.036 g/mL - Iwaniuk & Nelson, 2002), we used brain volume and mass interchangeably (compare Hurlburt et al., 2013; Herculano-Houzel, 2023). For maniraptoriform species, because their endocasts preserve brain contours similar to those of avians, we assumed a brain:endocast ratio of 100%. This is consistent with empirical data on the relationship between MBr and EV in modern birds, which suggest contributions of meningeal tissue to endocast volume to be negligible (Iwaniuk & Nelson, 2002). For other dinosaurs, we assumed MBr:EV ratios of 31% and 42%. Many previous studies have assumed a 50% ratio in these groups (reviewed in Morhardt, 2016) while some even assumed 100% (Balanoff et al., 2013) or advocated for intermediate values (e.g., Evans et al., 2008; Knoll & Schwarz-Wings, 2009; Knoll et al., 2021). The widely adopted 50% ratio was originally proposed by Jerison (1973) and based on measurements from a likely subadult green iguana (*Iguana iguana*) and a mere visual estimate of endocranial filling in the tuatara (*Sphenodon punctatus*; the endocranial fill in this species is now known to be 30% in adults - Roese-Miron et al., 2023). We abandon the problematic 50% estimate and replace it here by the two aforementioned ratios that are based on the morphology of extant crocodilians, the closest extant analogs to most non-avian dinosaurs in regards to endocranial tissue organization, body size and braincase ossification. Excluding one anomalous value, the lowest MBr:EV ratio among the three longest American alligators (*Alligator mississippiensis*) studied by Hurlburt et al (2013) was found to be 31% (n_total_= 12, note that this figure excludes the olfactory bulb and tract portions of the endocranial cavity). This is consistent with observations on the largest Nile crocodile (*Crocodylus niloticus;* a 16-year-old female) studied by Jirak & Janacek (2017) when excluding the olfactory tracts and bulb portion of the endocast. The 42% ratio is derived solely from American alligators. An endocranial fill of 42% was found in an adult female with a total length of 2.87 m, which roughly approximates both (*a*) the maximum length for a female American alligator and (*b*) the midpoint length within the size spectrum of sexually mature alligators (Woodward et al, 1991; Hurlburt & Waldorf, 2002; Hurlburt et al, 2013).

The majority of EV data for Mesozoic dinosaurs were taken and modified from the literature (detailed out in Tab. 1). In many cases, original sources communicated measurements that correspond to total EV. This is the volume of the entire endocast, often including the region of the olfactory tract and bulbs as well as portions of the cranial spinal cord among other structures. For our analysis, we exclusively relied on the so-called “brain” endocast volume instead (BrEV; Fig. 3), which was popularized by Jerison (1973) and has been commonly used since then (e.g., Hurlburt, 1996; Larsson et al., 2000; Paulina-Carabajal & Canale, 2010; Hurlburt et al., 2013). It excludes the spinal cord portion of the endocast caudal to cranial nerve XII, the volume of nerve trunks from infillings of respective foramina and blood vessel casts, the labyrinth of the inner ear, the infundibulum, the pituitary fossa, and especially the volume of the olfactory bulbs and tracts (Fig. 4). The latter are often only poorly preserved in fossil endocasts, so that relying on specimens with intact casts of olfactory structures would have reduced our sample size.

**Figure 3.**
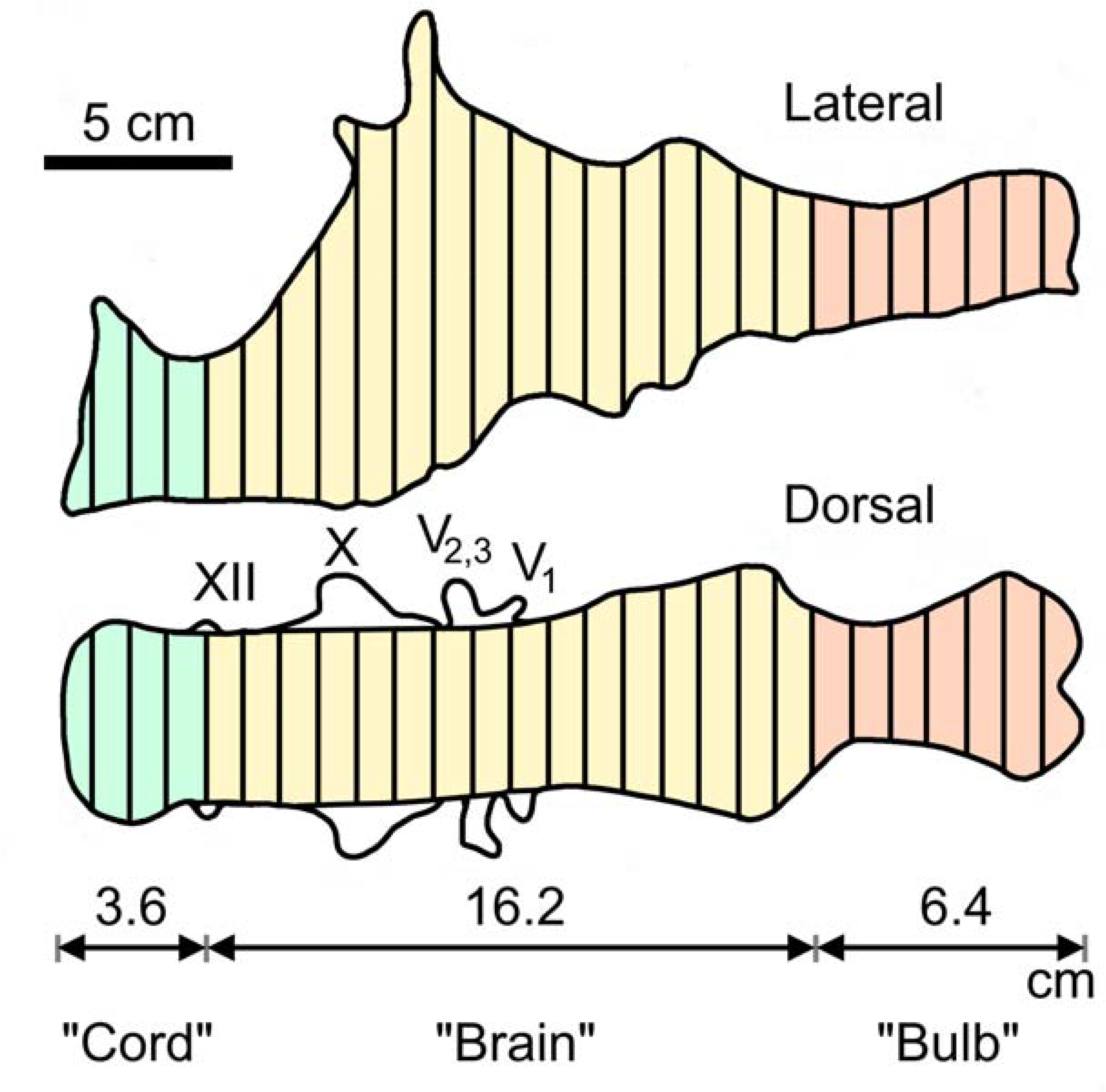
Exemplary graphic double integration (GDI) of the endocast of *Tyrannosaurus rex* AMNH 5029. Equally spaced lines are drawn across the right lateral and dorsal views respectively. Mean lengths of the lines drawn across the “brain” portion (BrEV) were 4.8 cm and 6.6 cm for dorsal and lateral views respectively. BrEV = π x 0.25 x 4.8 x 6.6 x 16.2 = 404 mL (the volume of the entire endocast was 536 mL). Abbreviations: “Bulb”: Olfactory tract and bulbs; “Cord”: spinal cord; V: Trigeminal nerve with its three branches (V1, V2,& V3); X: vagus nerve; XII: hypoglossal nerve. (Adapted from Fig. 2.7 in Jerison,1973, p. 51).

**Figure 4:**
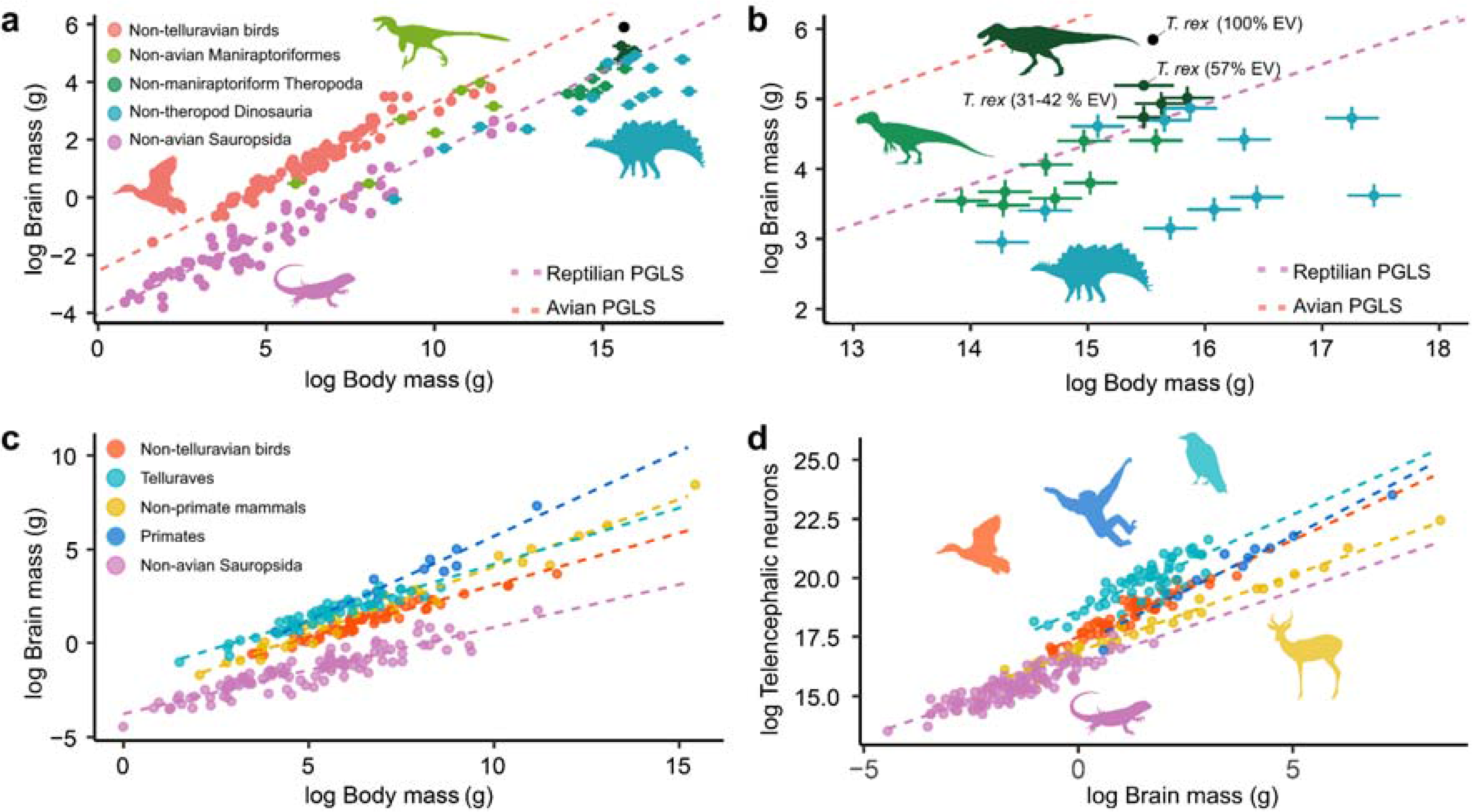
Relative brain size and forebrain neuronal numbers in Mesozoic dinosaurs and other amniotes. **a**.The log-transformed mass of the brain is plotted as a function of the mass of the body for extant and fossil sauropsids. In the case of fossil species, the mean body and/or brain size is shown along with standard deviations. The orange dotted line represents the regression line for avian species (excluding the large-brained clade Telluraves) obtained from PGLS while the pink one represents the same for extant non-avian sauropsids (“reptiles’’ in the colloquial sense). **b.** A detail of the plot shown in **a**. to illustrate the range of relative brain size in *Tyrannosaurus rex* and other Mesozoic dinosaurs that we consider plausible. Besides our own brain size estimates, the plot contains those from Morhardt (2016) (specimen AMNH FR 5117, endocranial fill = 57%) and Balanoff et al. (2013) (specimen AMNH 5029, endocranial fill = 100%, assumed MBd = 5840 kg) **c.** Plot showing log-transformed brain mass for different groups of extant amniotes plotted against body mass. **d.** Plot showing log-transformed numbers of telencephalic neurons as a function of the mass of the forebrain, illustrating neuronal density. Note that non-avian sauropsids and non-primate mammals differ only moderately from one another here, although mammals have markedly larger brains relative to body size, as shown in **c**. See methods for data sources. Silhouettes were taken from PhyloPic (listed clockwise from top left): *Anas* (in public domain) *Morunasaurus* (in public domain), *Dromaeosaurus* (in public domain), *Stegosaurus* (by Matt Dempsey), *Allosaurus* (by Tasman Dixon), *Tyrannosaurus* (by Matt Dempsey), *Corvus* (in public domain), *Hylobates* (by Kai Caspar), *Antidorcas* (by Sarah Wernig).

In some dinosaurs, there is an obvious constriction and/or a change in surface morphology at the junction of the cerebrum and olfactory tract (e.g., *Euoplocephalus tutus* AMNH 5337 - Hopson, 1979; *Stegosaurus ungulatus* CM 106 - Galton, 2001; *Diplodocus longus* CM 11161- Witmer et al., 2008). If present, this was used as a landmark to delineate these brain regions from one another. In less obvious cases, the junction between the cerebrum and olfactory tract portion was assumed to be where the ventral curve of the rostral cerebrum flattens out to approach a horizontal orientation. When selecting the boundary, we erred towards a more rostral location, to assign as much of the endocast as part of the BrEV as seemed reasonable. In American alligators, the rostral termination of the cerebrum within the rostral subarachnoid space is clearly visible (Fig. 2C; SaSR) and consistent with the change in curvature referred to above.

We used manual GDI to extract BrEV from total EV (relevant details for each specimen are provided in Supplementary File 1 Part B). The method involves drawing an outline around two scaled orthogonal two-dimensional views of an endocast, and adding equally spaced lines perpendicular to the endocast midline (Fig. 3). The mean length (cm) of these lines in each view (i.e., dorsal, lateral) provides diameters D1 and D2. The volume (mL) of the desired region is calculated using these two diameters and the length (L, cm) in the formula for the volume (mL or cm^3^) of a cylinder where (all lengths in cm):

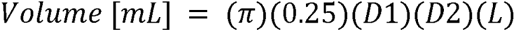

GDI has been demonstrated to produce reasonable estimates of endocast volumes (Fig. 3). For instance, Jerison (1973) used GDI to calculate a total endocast volume of 536 ml for a *T. rex* specimen (AMNH 5029), which was 101.13% of the 530 ml volume determined for it by means of water displacement (Osborn, 1912; Fig. 3). For the same specimen, Jerison (1973) calculated a 404 ml volume for the “brain region” of the endocast (extending from cranial nerve XII to the rostral cerebral limit), which was 106.04 % of the 381 ml obtained by CT volumetry (Hurlburt et al., 2013).

### Body mass estimates

We calculated the body mass (MBd, g) of the selected dinosaur taxa (and its mean absolute percent prediction error - PPE or %PE - Campione & Evans, 2020) in a standardized manner based on the minimum femoral circumference (as well as humeral circumference in case of quadrupedal taxa) by aid of the *QE()* and *cQE()* functions from the MASSTIMATE package (Campione, 2020) in R (R Core Team, 2023). Data on relevant long bone dimensions were primarily obtained from Benson et al. (2017). Corresponding specimens as well as additional sources and information on stylopodium circumference measurements are listed in Tab. 1. To the best of our knowledge, all data correspond to adult specimens.

### Phylogenetic modeling of neurological variables

We used data from extant sauropsids to place brain size variables for Mesozoic dinosaurs into their phylogenetic context. To examine variations in the relative size of the brain in fossil taxa and to calculate potential neuronal scaling regimes in extinct dinosaurs, we relied on log-transformed published data on brain mass, telencephalon mass and telencephalic neuronal numbers in extant groups (see below). Allometric equations were calculated with least squares linear regressions using phylogenetic generalized least squares (PGLS) to account for phylogenetic relatedness (Garland and Ives, 2000). PGLS allows the covariance matrix to be modified to accommodate the degree to which trait evolution deviates from Brownian motion, through a measure of phylogenetic correlation, Pagel’s λ (Pagel, 1999). PGLS and maximum likelihood estimates of λ were performed using the *ape* (Paradis & Schliep, 2019) and *nlme* (Pinheiro et al., 2017) packages in R.

To compare differences in relative brain size across groups, phylogenetically corrected ANCOVA with Tukey post hoc comparisons were performed using a modified version of the *multcomp* package (Hothorn et al., 2015; modification allowed outputs of the *nlme* package (*gls* objects) to be processed). Because of the uncertainty in estimating both brain mass and body mass in Mesozoic dinosaurs, we opted to test for inter-group differences in two datasets: One with the greatest possible relative brain size i.e., the lowest body mass estimate (lower PPE) for each species and the brain mass estimated from the highest assumed endocranial fill (42%), and one with the lowest relative brain size i.e. the highest body mass estimates (upper PPE) and the lowest assumed endocranial fill (31%). Since we assumed that the brain filled 100% of the endocranial cavity in maniraptoriform theropods, inferred relative brain mass for these species was only affected by differences in the applied body mass estimates.

As mentioned above, an important assumption of Herculano-Houzel (2023) is that theropods in general had relative brain sizes similar to birds. However, there is notable discontinuity in relative brain size and brain morphology between maniraptoriforms and more basal non-maniraptoriform theropods (Fig. 1, Balanoff et al., 2013). Because of this, we divided our sample of Mesozoic theropods into these two groups (Tabs. 1 & 2). For *T. rex*, mean values for the three available adult brain mass and corresponding body mass estimates were used. We grouped Sauropodomorpha and Ornithischia together as non-theropod dinosaurs and compared relative brain size in this group with that in the two theropod samples. PGLS models of brain mass vs. body mass with clade as a covariate were used to test if relative brain size in these groups differs significantly between them and from extant birds and/or non-avian sauropsids. Relative brain size data for 63 extant non-avian sauropsids (including lepidosaurs, crocodilians, and turtles) and 84 bird species (not including members of the large-brained clade Telluraves) were derived from Hurlburt (1996), Chentanez et al. (1983) and Roese-Miron et al. (2023) and are listed in Supplementary File 2. Importantly, these sources provide brain mass estimates excluding the olfactory tracts and bulbs and thus fit our brain size estimates for Mesozoic dinosaurs.

To test for differences in relative brain size we built a phylogenetic tree for all 175 fossil and extant species. To construct the phylogeny of bird species, we extracted 1,000 fully resolved trees from birdtree.org (Jetz et al., 2012) using the Hackett et al. (2008) backbone, and built a maximum clade credibility (MCC) tree using *phangorn* (Schliep, 2011). For lepidodaurs, we followed Kverková et al. (2022) by using a species level time-calibrated phylogeny (Tonini et al., 2016) and built a MCC tree the same way as we did for birds. For phylogenetic information on turtles and crocodilians, we relied on the Timetree of Life (Kumar et al., 2017). We then stitched the trees together manually, using the divergence times from the Timetree of Life. For Mesozoic dinosaurs (31 species) we used an updated version of the composite phylogeny of Benson et al. (2014, 2018). Phylogenies for fossil dinosaurs, extant non-avian sauropsids, and birds were stitched together manually using Mesquite (Maddison & Maddison, 2023). We opted to set all branch lengths to 1. This was done because clade-specific trees were obtained from various sources applying different phylogenetic methods, which, together with issues related to the precise dating of some of the fossils covered, makes it difficult to have well calibrated branch lengths. Importantly, simulation studies have found that independent contrasts and PGLS are robust to errors in both phylogenetic topology and branch lengths, so that we do not expect uniform branch lengths to compromise our analyses (Díaz-Uriarte & Garland 1998; Symonds 2002; Martins & Housworth 2002; Stone 2011).

Tree building procedures were the same for telencephalic neuron count regressions, but trees used here included branch lengths. We calculate regression lines between brain mass and telencephalic number of neurons for extant non-telluravian birds and non-avian sauropsids. Analogous to Herculano-Houzel (2023), these regressions were then used to estimate telencephalic neuron counts in dinosaurs, applying either an avian or a reptilian scaling regime. Since our estimates are based on brain portion endocasts that exclude the olfactory system, our telencephalic neuron counts correspond to the pallium and subpallium. Data on whole brain and telencephalic neuron counts as well as on total telencephalic and brain mass (including olfactory tract and bulbs, since neuron count data excluding these structures are currently unavailable for the taxa considered here) for birds (*n* = 112) were derived from Kverková et al. (2022) and Sol et al. (2022), for non-avian sauropsids (*n* = 108) from Kverková et al. (2022) and for mammals (*n* = 39) from Herculano-Houzel et al. (2015). The dataset is included in Supplementary File 3.

To get a more precise estimate of the possible number of telencephalic neurons in *T. rex*, we also modeled scaling regimes for telencephalon mass vs. telencephalic neuron numbers in extant sauropsids (non-avian sauropsids and non-telluravian birds), using the same references listed above. We then calculated telencephalic neuron numbers in *T. rex* using the obtained scaling regimes and applying the telencephalic volumes estimated with a comparative 3D landmark approach by Morhardt (2016) (referred to as “cerebral hemispheres” therein, excluding olfactory bulbs and tracts) for specimen AMNH FR 5117. Based on the estimates of Morhardt (2016), we also comparatively assessed the mass of the telencephalon and cerebellum in *T. rex*.

We want to note that our neuron count estimates might be biased by the fact that we predict neuron numbers in the pallium and subpallium (telencephalon excluding the olfactory system) based on total telencephalic neuron counts (including the olfactory system) here. This is an issue that in parts also applies to Herculano-Houzel (2023) and which we cannot circumvent due to limitations of the available raw data.

## Results

### Relative brain size

We did not recover notably large relative brain sizes in large-bodied theropods like *T. rex*. Instead, our analyses suggest that these animals had relative brain dimensions comparable to extant non-avian sauropsids such as lizards and crocodilians, as did Mesozoic dinosaurs outside of the clade Theropoda (Table 3). Relative brain sizes similar to those of extant birds appear to only have emerged among the maniraptoriform theropods: PGLS models showed a significant difference in relative brain size (intercept) between non-maniraptoriform theropods, such as *T. rex*, and the more bird-like Maniraptoriformes, which tended to have larger brains than other dinosaurs (PGLS, max: *F*_4,172_= 9.49, *p* ≤ 0.0001, λ = 0.707); min: *F*_4,172_= 14.03, *p* ≤ 0.0001, λ = 0.707; Fig. 4a-b). Post-hoc analysis shows that both the maximum and minimum relative brain size estimates for non-maniraptoriform theropods like *T. rex* are not significantly different from what would be expected from extant non-avian sauropsids (Tab. 3). However, both minimum and maximum relative brain size estimates for these animals are significantly smaller than what would be expected for extant birds (Tab. 3; Fig. 4). On the other hand, we found that maniraptoriforms show no significant differences in relative brain size compared to birds (Tab. 3) regardless of whether maximum or minimum relative brain size was assumed (Tab. 3; Fig 4; note that some maniraptoriforms such as *Shuvuuia deserti* cluster with non-avian sauropsids rather than with birds, though). In contrast to maniraptoriforms, other theropods did not exhibit significantly larger brains than the non-theropod dinosaurs of the clades Sauropodomorpha and Ornithischia (Tab. 3), data for which we pool here. Relative brain sizes in these dinosaurs were not recovered to differ notably from those of non-avian sauropsids. However, if minimum figures are assumed, their relative brain sizes would have been notably small, approaching a significant difference to extant non-avian sauropsids (Tab. 3).

### Numbers of neurons

We re-calculated estimates for the number of forebrain neurons in Mesozoic dinosaurs based on PGLS-derived regressions of brain size vs. number of telencephalic neurons in extant non-avian sauropsids and birds. Our neuron count estimates are listed in Table 2 and are compared to those of Herculano-Houzel (2023), whereas regression parameters are provided in Table 4. While many of the estimates do not differ notably from one another, the differences for some taxa, especially large theropods, are striking. For *T. rex*, Herculano-Houzel (2023) provided an estimate of 300-450 M forebrain neurons if modeled based on extant non-avian sauropsids, and 2-3B based on an avian regression. In contrast, we estimated a range of 245-360M neurons with a reptilian regression and ∼1-2 B with an avian one (Tab. 2, Fig. 5). Using the forebrain volumes estimated for *T. rex* by Morhardt (2016), we predict 133-166 M telencephalic neurons in this species if applying a reptilian scaling and 0.989 to 1.25 B based on an avian scaling (Fig. 5).

**Figure 5:**
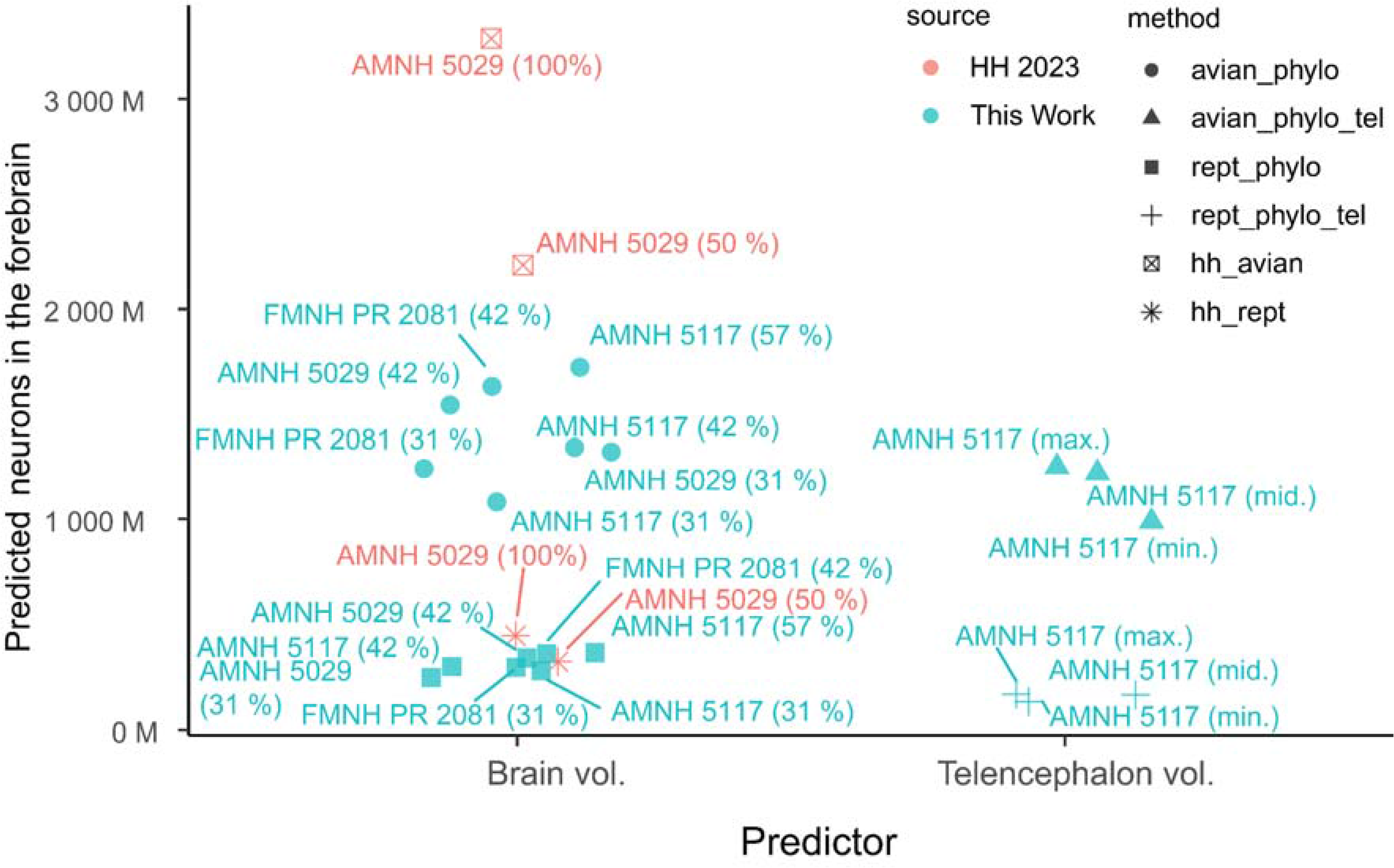
Predicted numbers of neurons in telencephalon (excluding olfactory tracts and bulbs) of *Tyrannosaurus rex*. Points represent the estimated number of neurons in three adult specimens of *T. rex* using different inference methods. Estimates from this study using a regression that takes phylogenetic relationships into account (PGLS, see methods, filled circle, triangle, square and cross) are plotted in cyan. The estimates from Herculano-Houzel (2023) based on a non-phylogenetic regression are shown in red (crossed square, asterisk). Different underlying ratios of brain volume:endocranial volume are annotated. On the left predicted numbers of forebrain neurons (based on either the avian or extant non-avian sauropsid scaling regime) based on the estimated volume of the brain portion of the endocast are shown. On the right, analogous to that, the predicted numbers of telencephalic neurons based on forebrain volumetric estimates by Morhardt (2016) is plotted.

**Table 2:**
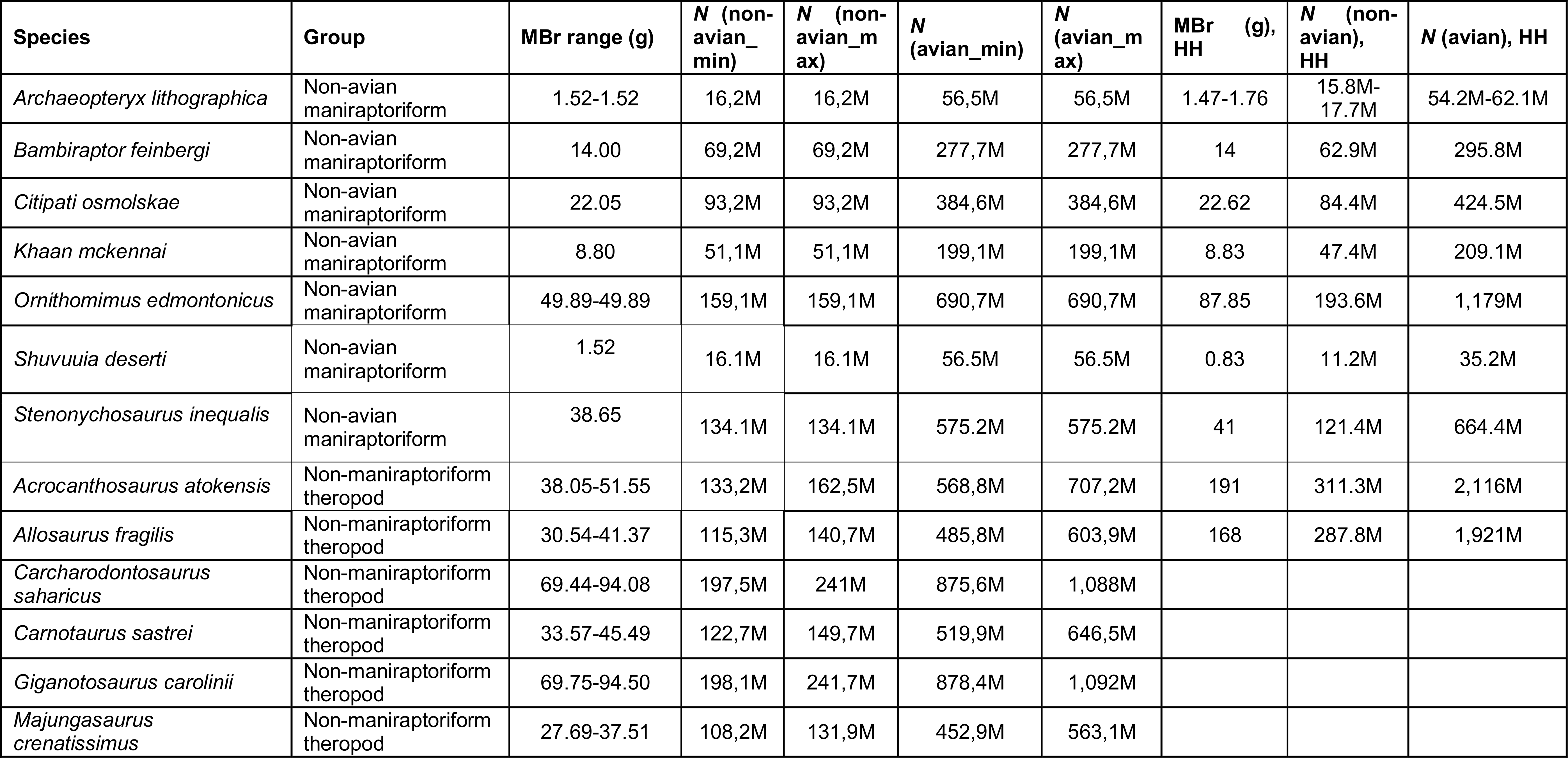
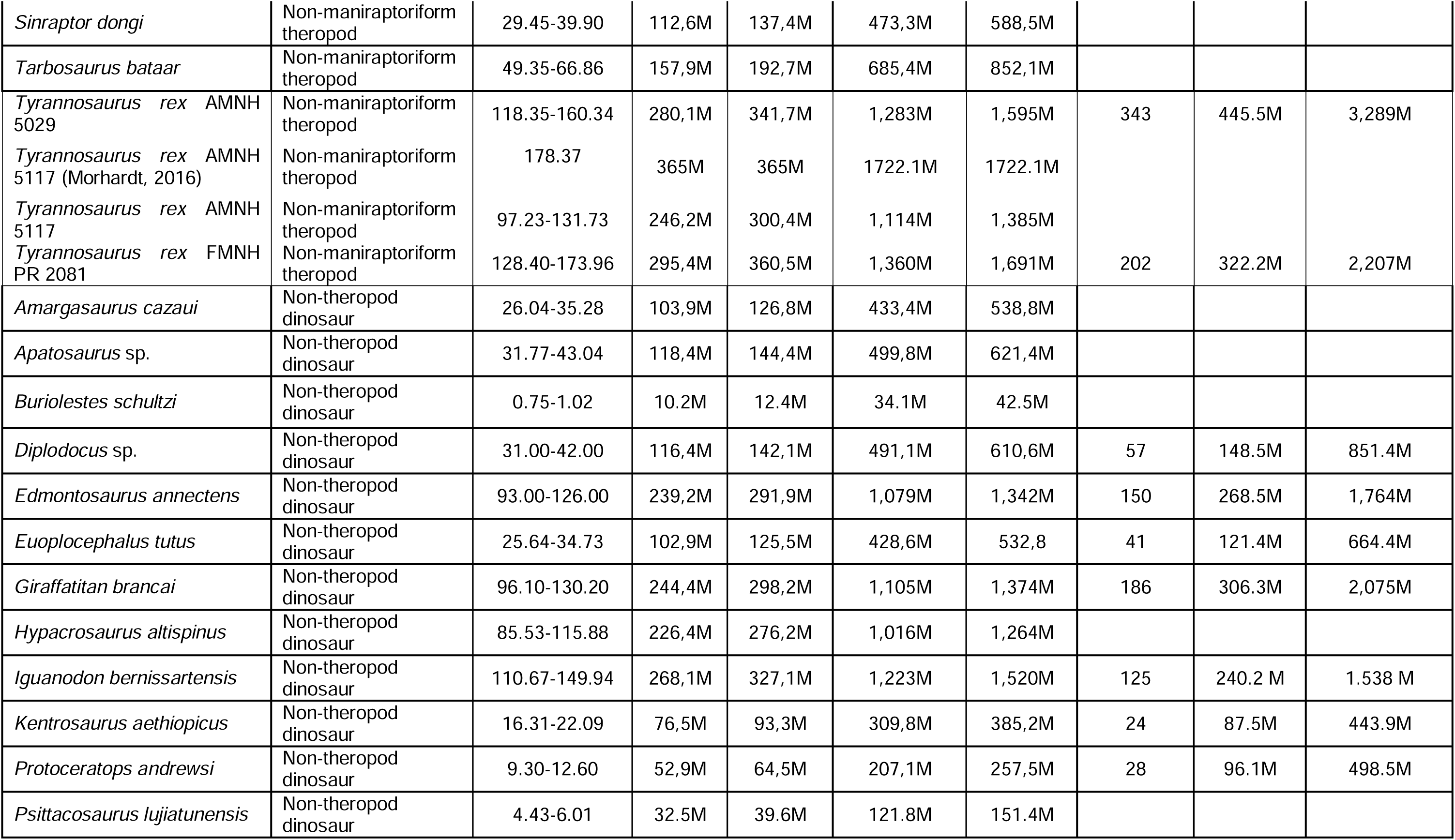
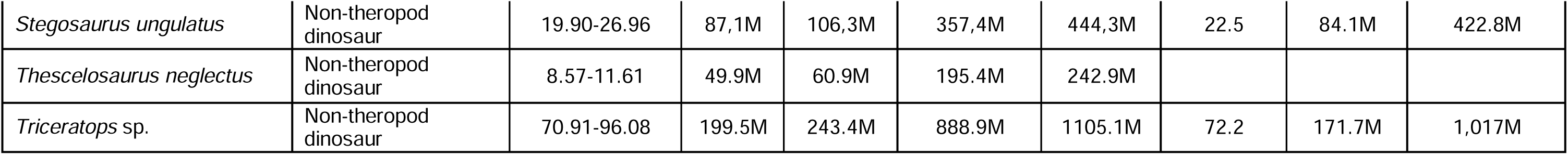
Estimates of telencephalic neuron counts (*N*; excluding the olfactory system) in Mesozoic dinosaurs. Our inferences are compared with those presented by Herculano-Houzel (2023; HH) if respected species were included in both studies (see text for the rationale of our sample composition). Minimum and maximum estimates based on both avian and non-avian sauropsid regressions are provided.

**Table 3:**
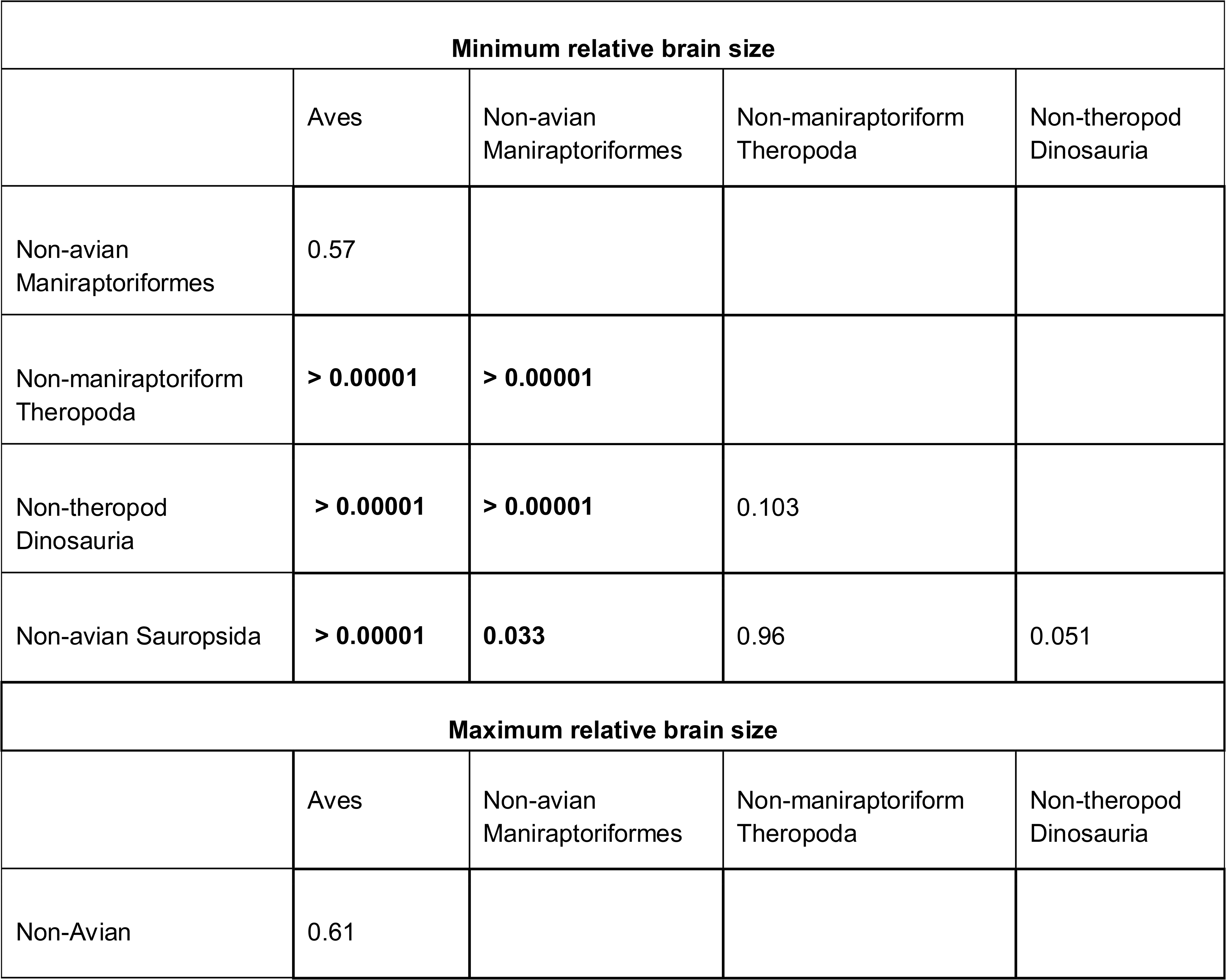
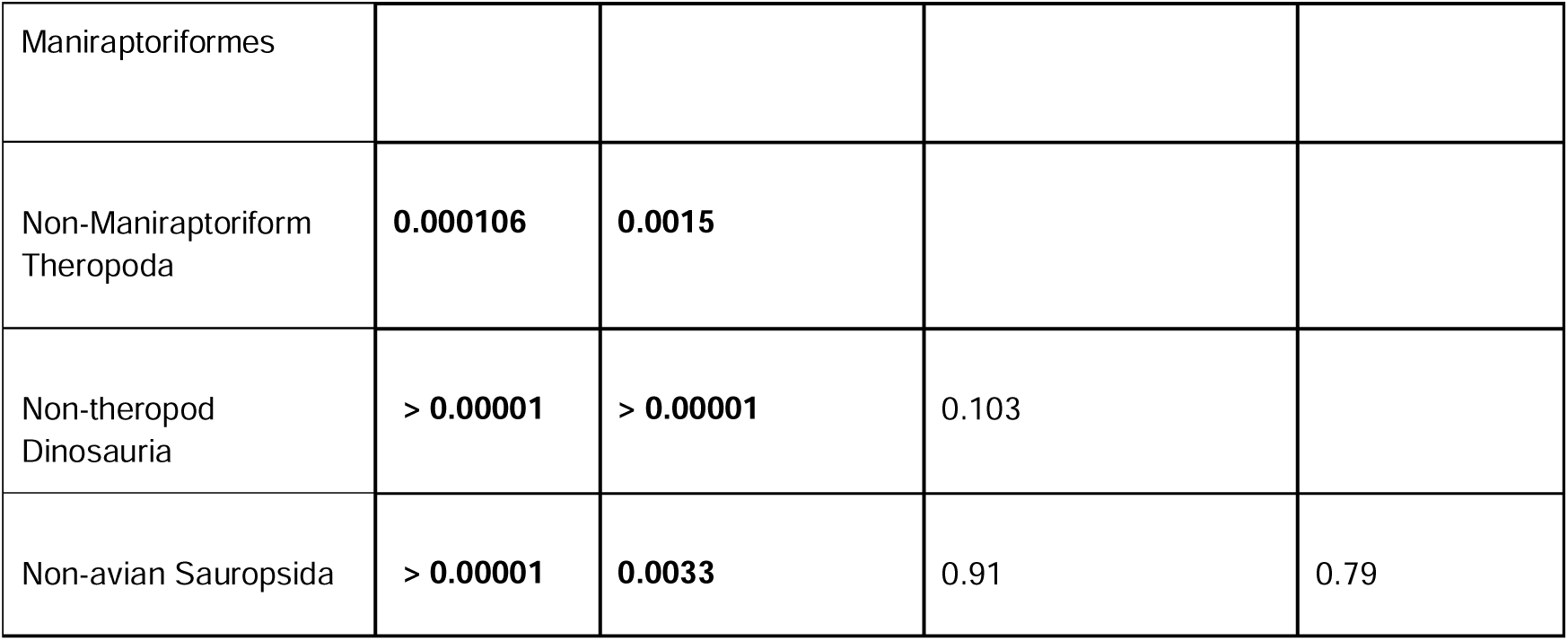
Tukey post hoc comparisons for a phylogenetic ANCOVA testing for differences in relative brain size between groups of fossil dinosaurs and extant sauropsids. Significant *p*-values are shown in bold.

**Table 4:**
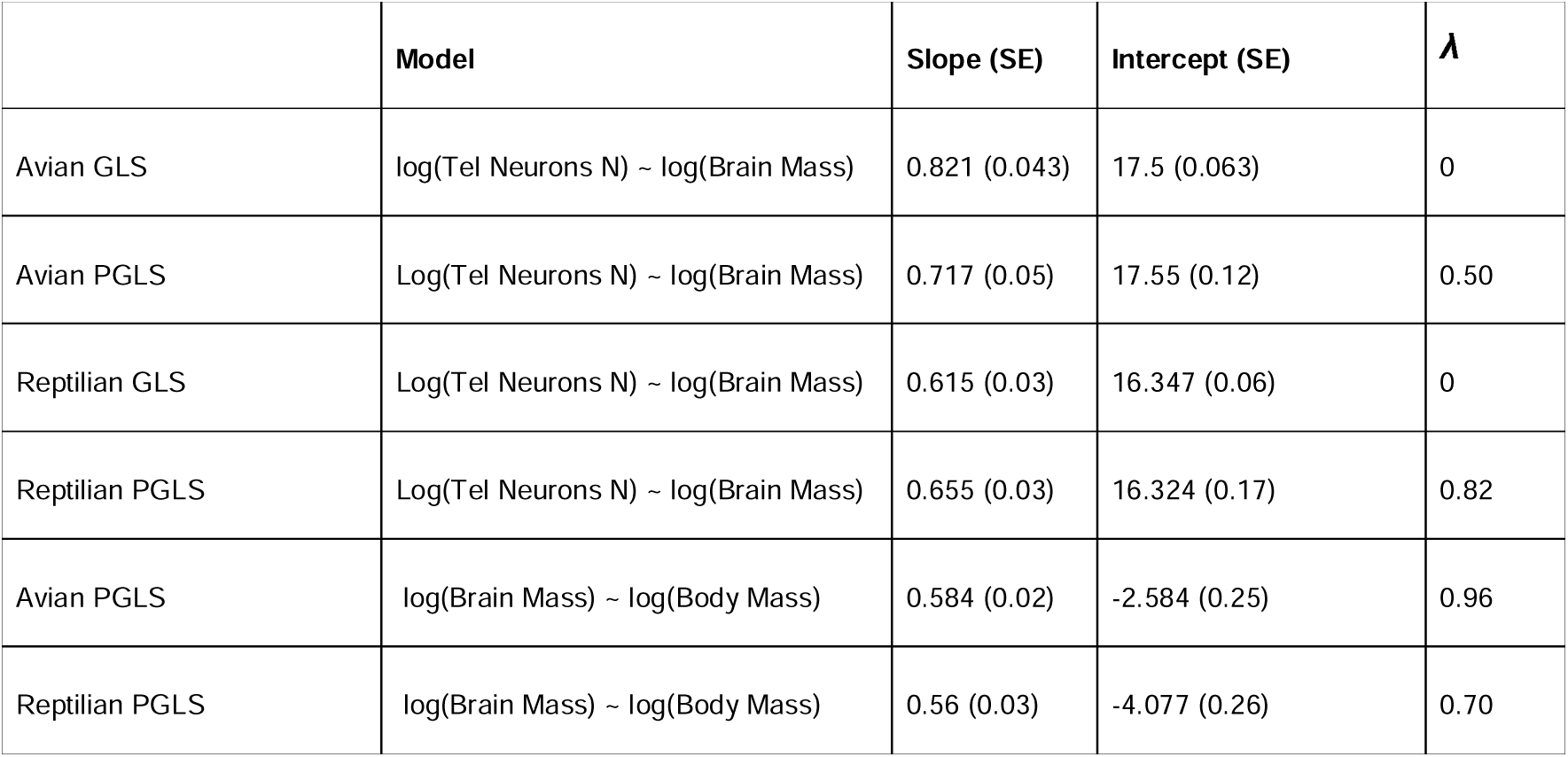
Regression parameters for different models describing the scaling of neurological traits in extant non-avian sauropsids (“reptiles”) and non-telluravian birds. Pagel’s λ (ranging between 0 - 1) was used to quantify the phylogenetic signal. See methods for details. SE = standard error.

## Discussion of empirical results

We want to emphasize two aspects of our empirical findings that contrast with those of Herculano-Houzel (2023). Firstly, we did not find relative brain size to separate non-maniraptoriform theropods such as *T. rex* from extant non-avian sauropsids like crocodilians and lizards or other large-bodied dinosaurs outside the clade Theropoda; rather, our data support a grade shift in this trait between maniraptoriform and non-maniraptoriform theropods, which at least in parts relates to an increase in endocranial fill. As we have argued beforehand, we see no support for the brains of non-maniraptoriform theropods, sauropodomorphs, and most ornithischians to have filled the endocranial cavity in a bird-like fashion. However, we are aware that such a condition, or one that is at least intermediate between modern birds and crocodilians has been proposed for all of these groups at one point (e.g., Knoll & Schwarz-Wings, 2009; Morhardt, 2016; Balanoff et al., 2013; Knoll et al., 2021; see Supplementary File 1 Part C for further comments on that topic). Obviously, future research might significantly change our understanding of endocranial tissue organization in Mesozoic dinosaurs and thus challenge the assumptions that we make here. For the time being, however, we consider our crocodilian-based inferences more plausible and parsimonious than the alternative suggestions proposed so far.

Our approach suggests that relative brain size in all dinosaurs, except for the majority of maniraptoriform theropods, does not differ significantly from values present in extant non-avian reptiles. These results agree with previous conclusions (Hurlburt et al., 2013; Morhardt, 2016). Nonetheless, we want to stress that it remains unclear how meaningful the transfer of brain size scaling rules established for the given extant bird (32g – 120kg) and non-avian sauropsid (1g – 71kg) datasets to large-bodied dinosaurs actually is. Brain-body size ratios in extant cetaceans drop dramatically in taxa that evolved multi-ton body masses (Tartarelli & Bisconti, 2006), suggesting that allometric trajectories need to be accounted for. However, the restricted body mass spectrum of extant birds and reptiles as well as the limited availability of large-bodied crocodilians and turtles for neurological research hinders the compilation of such datasets for sauropsids. Furthermore, the scarcity of complete and adult dinosaur endocasts from taxa that also preserve stylopodial elements to derive body mass estimates from, limits our understanding of differences in brain size scaling between taxa. Different clades of mammals and birds have been shown to have distinct allometric relationships for relative brain size (Ksepka et al., 2020; Smaers et al., 2021). The same might have been the case in non-avian dinosaurs, biasing comparisons between the groupings we selected here. In addition to that, there might also be temporal effects on relative brain size. Such a phenomenon appears to be rampant in mammalian evolution during the Cenozoic (Bertrand et al., 2022). To our knowledge, this pattern has not been properly described yet in other vertebrate groups but should be considered in future studies on brain evolution in long-lived clades such as dinosaurs.

Secondly, our empirical findings do not support Herculano-Houzel’s (2023) claim of exceptionally high telencephalic neuron counts in dinosaurs, particularly in *T. rex* and other large theropods. Instead, *T. rex* likely did not exhibit more than approximately 1.5 B (or at a maximum 2 B) telencephalic neurons, even when an avian neuronal density is assumed. If we assume reptilian neuronal densities, it might even have exhibited neuron numbers an order of magnitude lower than the 3.3 B suggested by Herculano-Houzel (2023). Apart from the difficulty of estimating brain mass from a dinosaurian endocast, there is one additional caveat to our neuron count estimates that needs to be acknowledged and that also applies to Herculano-Houzel’s (2023) study: Telencephalic neuron numbers can only be reliably derived from total brain mass when the proportions of the studied brains are comparable. Since brain morphology in many dinosaurian lineages differs significantly from both extant birds and non-avian sauropsids (Fig. 1, Paulina-Carabajal et al., 2023), biases are thus ingrained into such estimates. Volumetric modeling of brain regions from endocasts, on which we relied here for *T. rex* exclusively, could potentially ameliorate this problem to some degree (Morhardt, 2016) but it is challenging and not yet widely used. For *T. rex*, such inferences yield lower telencephalic neuron numbers than would be hypothesized based on our total brain volume estimates, if reptilian scaling rules are applied (Fig. 5). They overlap if an avian neuron count scaling is assumed (Fig. 5).

We want to emphasize that there is little reason to assume that the brains of non-maniraptoriform theropods such as *T. rex* had a telencephalic neuronal density similar to that of extant birds. In living sauropsids, relative brain size is positively correlated with neural density (Kverková et al., 2022). We show that this measure likely did not differ significantly between those theropods and extant non-avian sauropsids. Consequently, relative brain size cannot be used as an argument to defend elevated neuron densities in these animals. The presence of endothermy in dinosaurs (see below) does also not entail avian neuronal density (contra Herculano-Houzel, 2023): Similar to birds, mammals have evolved endothermy and exhibit large relative brain sizes (Fig. 4C; Tsuboi et al., 2018). Furthermore, they display a unique multilayered cerebral neuroarchitecture (Briscoe & Ragsdale, 2018). Yet their average forebrain neuronal density is only moderately elevated compared to extant non-avian sauropsids (at least if anthropoid primates are not considered) and there is a broad overlap in neuronal density between the groups (Fig. 4D; Kverková et al., 2022), indicating remarkable conservatism in this trait. With respect to birds, however, the typical mammalian telencephalic neuron density is remarkably low (Fig. 4D). Interestingly, brain cell densities (suggestive of high neuron counts but including endothelial and glia cells) on par with or even higher than those of telluravian birds have recently been identified among ectotherm teleost fish, with comparatively small relative brain sizes (Estienne et al., 2024). All of this suggests that metabolic rate and neuronal density are not tightly coupled and that endothermy cannot be used as a proxy for the latter. Finally, the shape of the endocast and volumetric estimates of its forebrain and cerebellar portions (compare Fig. 6) suggest that the brains of *T. rex* and other large non-maniraptoriform theropods were not dissimilar to those of extant crocodilians (Rogers, 1998; Hurlburt et al., 2013; Morhardt, 2016), which reflect the plesiomorphic archosaurian condition (Fabbri & Bhullar, 2022). Given this morphological conservatism and the rather static neuron densities of non-avian amniotes, it appears appropriate to assume reptilian neuronal densities for these animals.

**Figure 6:**
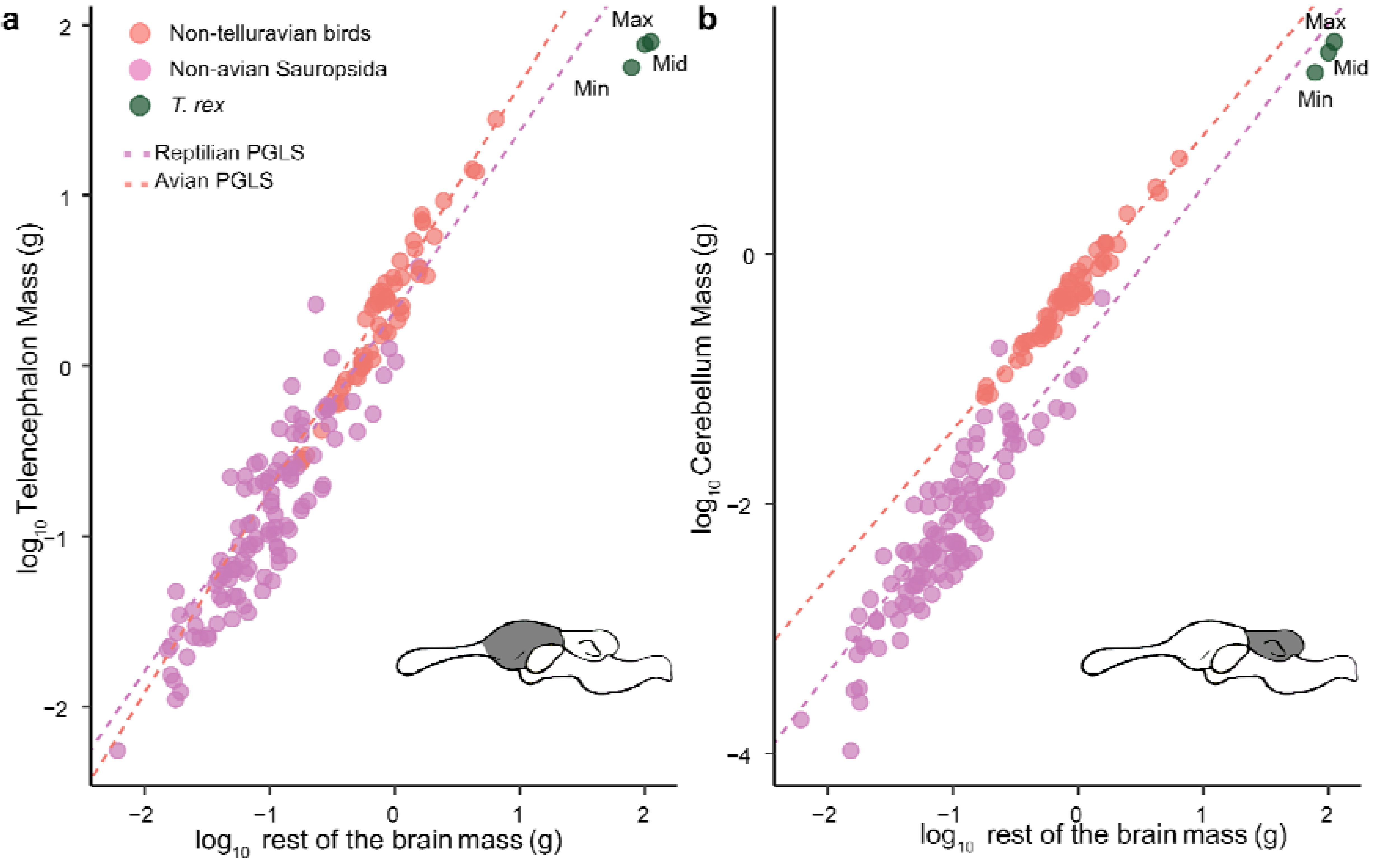
Relative size of telencephalon (excluding olfactory bulb and tracts) and cerebellum in T. rex. **a:** The log-transformed mass of the telencephalon in extant non-telluravian birds and non-avian sauropsids is plotted as a function of the mass of the rest of the brain (total brain - telencephalon - cerebellum volume). Green dots show the maximum, mid and minimum estimates for the mass of the *T. rex* telencephalon as estimated from digital endocasts of AMNH 5117 by Morhardt (2016). The orange dotted line represents the regression line for non-telluravian bird species obtained from PGLS while the pink represents the same for non-avian sauropsids. **B**: Analogous plot to **a**, but for the cerebellum. Note that telencephalic mass in extant species includes the olfactory bulb and tracts.

It is tempting to speculate that the increased neuronal density that sets extant birds apart from other sauropsids and mammals coevolved with the marked changes in brain morphology and size that occurred in maniraptoriform theropods. If we indeed assume that an avian-like brain organization and high neuronal density emerged early within this clade’s history, it is plausible that the Mesozoic dinosaurs with the highest neuron counts, perhaps above the extant avian range, are represented by the largest-bodied taxa within this group (for which no complete endocasts are currently available). These include bizarre animals such as the immense ornithomimosaur *Deinocheirus mirificus* (∼ 7 t), the scythe-clawed *Therizinosaurus cheloniformis* (∼ 5 t) and the giant oviraptorosaur *Gigantoraptor erlianensis* (∼ 2 t). Alternatively, the emergence of volancy in small maniraptoriforms similar to *Archaeopteryx* might have driven the evolution of elevated neuron densities, since active flight likely imposes constraints on skull and brain size (Olkowicz et al., 2016; Shatkovska & Ghazali, 2021). However, the lack of reliable morphological markers to infer neuron density renders all these notions speculative. Such a vagueness is inherent to predictions about the biology of extinct taxa without close living relatives and obviously needs to be acknowledged. The main argument for assuming avian neuronal densities in any group of Mesozoic dinosaurs is that the emergence of this trait within the avian stem-lineage cannot be reliably dated and thus might have significantly preceded the origins of crown birds. Hence, both a non-avian sauropsid and an avian neuron density (as well as intermediate conditions) could principally be justified for dinosaurs, although we advocate for the former if taxa outside the Maniraptoriformes are concerned. Importantly, however, even if we had robust evidence for high neuron counts in Mesozoic dinosaurs, this would by no means automatically suggest exceptional cognitive capacities.

## General discussion - implications for neuron count and brain size estimates for vertebrate paleontology

### 1) Are neuron counts good predictors of cognitive performance?

To infer cognitive abilities in extinct animals from neuron count estimates for the whole brain or pallium, we first need to be assured that this measure can give us meaningful insight into behaviors of extant ones. However, while there is some evidence for effects of pallial neuron counts on species-level cognitive performance in primates (Deaner 2007 - but see below) and birds (impulse inhibition - Herculano-Houzel, 2017; but see Kabadayi et al. 2017 for conflicting evidence; foraging-related innovativeness – Sol et al., 2022; but consider limitations on how innovativeness is measured – Logan et al., 2018), the available data do not provide consistent support for the hypothesis that more neurons per se enhance cognition (Barron & Mourmourakis, 2023). As an example, Güntürkün et al. (2017) reviewed the performance of domestic pigeons (*Columba livia*), corvids and anthropoid primates in a number of cognitive tasks with the aim to determine if a “cognitive hierarchy” between the three groups exists. They note that pallial neuron counts in corvids are about 2–6 times lower than in large-bodied monkeys and apes but 6–17 times higher than in pigeons. Thus, one would predict major increases in cognitive capacities from pigeons to corvids to anthropoids. Yet, corvids typically perform on par with anthropoid primates (see also Kabadayi et al. 2016 and Pika et al. 2020), and pigeons do so as well in some cognitive dimensions, such as numerical competence and short-term memory (Güntürkün et al., 2017). In addition, standardized testing of various primate species suggests that small-brained lemurs with comparatively low neuronal densities (Kverková et al., 2022), monkeys and great apes rival each other in a number of cognitive dimensions (Schmitt et al., 2012; Fichtel et al., 2020). In fact, findings that report the influence of absolute brain size (and thus neuron number) on cognitive performance in primates (Deaner et al. 2007) have failed replication so far (Fichtel et al. 2020). As a final example, we want to point out large-bodied dolphin species, which have remarkably high neocortical neuron counts (*Globicephala melas* - 37 B, *Orcinus orca* - 43 B; Ridgway et al., 2019). Although neuron numbers in these animals vastly exceed those of humans (15-20 B), there is no evidence that cetacean cognition is on par or even superior to that of our species (e.g., Manger, 2013; Güntürkün, 2014). Hence, even immense differences in telencephalic neuron numbers do not necessarily create cognitive divides and their value in predicting cognitive performance is remarkably limited.

The case becomes even more untenable when we take more complex cognitive phenomena into consideration, such as habitual tool use. Remarkably, Herculano-Houzel (2023) suggested that this might be within the realm of possibility for large theropods such as *T. rex*, as it is for primates and telluravian birds. However, tool use even within these groups is rare, especially if the more rigorous definition of “tooling” (requiring the deliberate management of a mechanical interface, see Fragaszy & Mangalam, 2018) is employed: this occurs in only 9 avian and 20 primate genera (Colbourne et al., 2021). While it is true that telencephalon size in birds has an association with tool use (Lefebvre et al., 2002), this correlation does not hold any predictive power in the sense that all birds with a certain-sized telencephalon exhibit tool use. Even within corvids, which telencephalic neuron counts and sophisticated cognitive abilities overlap with those of anthropoid primates (Olkowicz et al., 2016; Ströckens et al., 2022), New Caledonian crows (*Corvus moneduloides*), and Hawaiian ‘alalā crows (*C. hawaiiensis*) are the only species known to employ and manufacture tools in the wild. Notably, both species inhabit remote islands, and they share unusually straight beaks and greater binocular overlap than other crows, which are thought to be specific morphological adaptations to enable tool use (Troscianko et al., 2012; Rutz et al., 2016). A similar situation can be observed in parrots. These birds display the highest avian telencephalic neuron counts (Olkowicz et al., 2016; Kverková et al., 2022; Ströckens et al., 2022), and a greatly enlarged medial spiriform nucleus, which acts as an interface between the pallium and the cerebellum, enabling enhanced motor cognition (Gutiérrez-Ibáñez et al., 2018). However, the Tanimbar corella (*Cacatua goffiniana*) is the only parrot known to be a sophisticated tool user in the wild (O’Hara et al., 2021); tellingly, the Tanimbar corella also inhabits an isolated Indonesian archipelago. Cases like these indicate that while there might be a chance that a gross neuron count threshold must be met for such sophisticated vertebrate tool use to emerge (a notion we would reject since ants evolved remarkable tool use skills with brains that are small and few in neurons even for the standard of hymenopteran insects - Godfrey et al., 2021), it is highly unlikely to happen without sufficient ecological pressure, and the differences between tool using and non-tool using species are likely too subtle to detect via measurement of neuronal quantities.

Considering these findings, it is unsurprising that taxa converging in neuronal counts often differ markedly in cognition and behavior. Herculano-Houzel (2023) ranked her neuronal count estimates for large theropods against those of anthropoid primates, but she might as well have done so for giraffes (1.7 B neurons), which exceed tool-proficient capuchins (1.1 B) and corvids (0.4-1.2 B) in telencephalic neuron numbers, rivaling macaques (0.8 - 1.7 B) (Olkowicz et al., 2016). We know little about giraffes’ cognitive abilities (Caicoya et al., 2018), but it would be appropriate to be skeptical of any claim that they might exhibit “macaque-like” cognition based simply on that measure. Too many other biological traits divide these taxa, perhaps most strikingly body size. While we agree with many contemporary authors that relative brain size per se is a flawed measure of cognitive complexity (e.g., Van Schaik et al., 2021), it must not be ignored. This is especially true if comparisons between primates and Mesozoic dinosaurs are drawn, since the species concerned may differ in body mass by several orders of magnitude. Contrary to the assumptions of Herculano-Houzel (2023), the size of the telencephalon and number of its neurons must be related to the dimensions of the body, because it processes sensory, visceral, and motoric information, which scale with body size (Chittka & Niven, 2009; Van Schaik et al., 2021). This fact is clearly reflected by the pronounced intra- as well as interspecific body size-dependent scaling of brain size in vertebrates (Tsuboi et al., 2018; Ksepka et al., 2020; Van Schaik et al., 2021; Bertrand et al., 2022), which can hardly be explained otherwise. Relative brain size and body size are thus not negligible variables in comparative cognition and need to be considered in paleoneurology.

The confounding factor of body size on neurological measures might be mitigated by calculating clade-specific portions of telencephalic mass dedicated to somatic functions (the regulation of visceral, sensory and motor processes unrelated to cognition) based on intraspecific variation (Triki et al., 2021; Van Schaik et al., 2021) or by focusing on neuron counts in brain regions that are evidently not involved in somatic processing (Herculano-Houzel, 2017; Logan et al., 2018). In fact, a number of studies, particularly in birds, were able to associate intraspecific differences in certain cognitive dimensions to localized neurological variation, making this approach a promising one (discussed by Logan et al., 2018). At the same time however, the great intra- and interspecific heterogeneity in brain tissue architecture and neurochemistry enormously complicates any interspecific extrapolations (Logan et al., 2018; Barron & Mourmourakis, 2023). Thus, researchers cannot translate these findings to extinct species with any tolerable degree of certainty. This issue is of special relevance when comparing sauropsids with mammals. The mammalian forebrain exhibits a layered cortex but the pallium of extant sauropsids (and thus likely Mesozoic dinosaurs) is largely nuclear in organization. As the forebrain increases in size and neuron counts, a cortical organization can reduce axon length (and therefore processing time and energetic demands) by bringing adjacent areas closer together through folding, something that is impossible in a nuclear organization (see Reiner, 2023 for an extensive review).

Neuron counts corresponding to major brain regions, whether empirically determined or estimated, dramatically simplify neuronal tissue complexity, as do measures such as absolute brain size or EQ. Based on current evidence, they also represent flawed cognitive proxies that need to be viewed in the broader context of an animal’s ecology, neuroanatomy, connectomics, and neurochemistry (Fields & Stevens-Graham, 2002; Eyal et al., 2016; Logan et al., 2018; Barron & Mourmourakis, 2023, Reiner, 2023). All in all, we want to discourage attempts to predict cognitive performance in extinct species based on endocast-derived neuron count estimates.

### 2) Inferring metabolic rate

Apart from inferences about cognition, Herculano-Houzel (2023) claims that relative brain size should be established as a new thermobiological indicator in vertebrate palaeontology: Relatively large brains, as inferred for theropods, should be viewed as indicators of endothermy, while smaller ones, as are found in pterosaurs, sauropodomorphs and many ornithischians, would indicate ectothermy. Whereas overall brain size in vertebrates is indeed correlated with metabolic rate (e.g., Yu et al., 2014 - but also note the extreme variability within ecto- as well as endothermic groups), Herculano-Houzel’s (2023) approach simplifies the matter and ignores a vast body of already available evidence on dinosaur thermobiology. First, as we have extensively discussed here, relative brain size in large theropods was probably markedly smaller than suggested by Herculano-Houzel (2023) and more similar to the condition in extant crocodilians rather than in birds. Second, it is important to point out that there is a spectrum of metabolic rates in vertebrates (Legendre & Davesne, 2020) rather than a dichotomy, as suggested by Herculano-Houzel (2023).

Where exactly certain ornithodiran taxa align within this spectrum continues to be debated, but there is consensus that dinosaurs and pterosaurs, despite their in parts iconically small brains, had metabolic rates well above the range of extant ectothermic sauropsids (see references below). Rather than emerging with theropods, contemporary evidence suggests that endothermy evolved in the ornithodiran stem-lineage or even earlier (Legendre et al., 2016; Benton, 2021; Grigg et al., 2022) and hence was inherited by pterosaurs and dinosaurs. The extensive data supporting the presence of endothermy across Ornithodira has recently been reviewed by Grigg et al. (2022) and includes the presence of hair-like, sometimes branched, integumentary structures (Benton et al., 2019; Campione et al., 2020), the efficiency of the ornithodiran respiratory system (Wedel, 2006; Butler et al., 2009; Aureliano et al., 2022; Wang et al., 2023), bone histology and high skeletal growth rates (de Ricqlès et al., 2000; Padian et al. 2004; Prondvai et al., 2012; Redelstroff et al. 2013; Legendre et al., 2016), paleoenvironmental data (Druckenmiller et al., 2021), models of locomotor costs (Pontzer et al., 2009) and geochemically-derived thermometric findings (Barrick et al., 1996; Dawson et al. 2020; Wiemann et al., 2022). Nevertheless, osteohistological evidence suggests that both theropod and non-theropod ornithodiran taxa varied in their growth and associated metabolic rates (Jenkins et al., 2001; Erickson et al., 2009; Redelstroff et al. 2013; D’Emic et al., 2023) and a secondary reduction of metabolic rate in some ornithischian groups appears plausible (Padian et al., 2004; Redelstroff and Sander, 2009; Wiemann et al., 2022), albeit still compatible with an endothermic physiology (Grigg et al., 2022).

Overall, we want to emphasize the need for a nuanced perspective on this trait. The assumption that relative brain size alone (even if inferred correctly) can outperform all the aforementioned thermophysiological predictors to infer endothermy appears at best improbable. Its utility to gauge metabolic rate across ornithodiran groups therefore remains highly doubtful and must be viewed in the framework of other, more robust lines of evidence.

### 3) Inferring life history traits

Finally, Herculano-Houzel (2023) suggests that neuron count estimates can be used to model life history traits in Mesozoic ornithodiran taxa. This notion is based on previous empirical work that showed an association between pallial neuron counts and selected ontogenetic variables in extant mammals and birds (Herculano-Houzel, 2019). Applied to *T. rex*, the respective equations predict that females reached sexual maturity at an age of 4–5 years and that the longevity of the species was 42-49 years (Herculano-Houzel, 2023).

These calculations rest on the assumption that *T. rex* had to have 2.2 – 3.3 billion pallial neurons. As we have shown, this premise appears exceedingly unlikely. Furthermore, the aforementioned life history predictions are contradicted by the fossil evidence: Sexual maturity in extinct nonavian dinosaurs can be estimated histologically by the presence of medullary bone, a tissue that forms as a calcium reservoir for egg shell production and which is also seen in female birds (Schweitzer et al., 2005; Woodward et al., 2020). The earliest estimate of sexual maturity in *T. rex*, as estimated by the presence of medullary bone, is 15 years (Woodward et al., 2020; Carr, 2020). If the life history of *T. rex* was similar to extant American alligators where sexual maturity occurs in animals that attain half of adult size (which would be in line with the available fossil data - Carr, 2020), then the earliest onset of sexual maturity in *T. rex* happened in its 12th year of life. Based on these lines of evidence, Herculano-Houzel’s (2023) method greatly underestimates the onset of sexual maturity by 8 to 11 years. Based on the number of lines of arrested growth in its long bones, which are thought to indicate annual cessations of growth, the chronologically oldest *T. rex* sampled so far lived up to 33 years (Cullen et al., 2020). Although it is not unreasonable to assume that *T. rex* lived longer than three decades, there is yet no histological evidence to support that hypothesis. Given that Herculano-Houzel’s (2023) longevity estimate is based on problematic premises, it should not be considered a plausible alternative.

In fact, if applied to species other than *T. rex*, the limitations of the aforementioned approach become even more visible. For instance, if the life history of the sauropod *Apatosaurus*, a gigantic dinosaur with an adult weight exceeding 30 tonnes, is modeled based on our own neuron count estimates derived from an avian regression and an endocranial fill of 42%, the equations suggest a longevity of only 24.5 years and an onset of sexual maturity at 2.2 years (note that assuming a non-avian sauropsid regression or smaller brain size would result in an even more fast-paced life history prediction). These figures are obviously unfeasible. We are aware that Herculano-Houzel (2023) assumes that sauropods such as *Apatosaurus* were ectothermic animals so that the given equations could not be applied to them. However, since this notion defies essentially all available evidence on the biology of sauropods (see above), we choose to ignore it here. To conclude, relationships between life history and neurology that were established from a selection of extant mammals and birds cannot be used to reliably infer ontogenetic parameters across non-avian dinosaurs (and potentially other fossil groups). We strongly discourage relying on them in palaeontological practice.

### Beyond endocasts: What are the limits of inference on dinosaur cognition?

If neuron count estimates and other endocast-derived variables do not allow reliable predictions about the cognitive abilities of non-avian dinosaurs to be made, what other methods are available? First of all, trace fossils can provide direct evidence on how dinosaurs exploited their environment and interacted with both hetero- and conspecifics (e.g., Carpenter et al. 2005; Varricchio et al. 2007; Lockley et al. 2016; Brown et al. 2021). While such fossils can provide precise and diverse insights into dinosaur behavior, obvious limitations render perspectives gained from them extremely patchy, nonetheless.

One further way of inferring cognitive traits in dinosaurs is by comparatively studying relevant behavioral phenomena in living crocodilians and birds, the groups that form their extant phylogenetic bracket. While such approaches are starting to gain pace (Zeiträg et al., 2023), we are not aware that ethological research could so far identify shared physical or social cognitive skills in crocodilians and birds that have not also been found in turtles and squamates (in case such comparative data is indeed available - Zeiträg et al., 2022; Font et al., 2023). Thus, the behavioral resolution of such approaches appears limited thus far. Cognitive traits identified exclusively in birds or crocodiles cannot simply be extrapolated to Mesozoic dinosaurs with any degree of certainty since they might represent crown group apomorphies. Whereas it might be appealing to hypothesize that cognitive patterns found among modern palaeognaths are representative for their maniraptoriform forerunners (Jensen et al., 2023; Zeiträg et al., 2023), this idea is (in most cases) not testable and should hence not be disseminated uncritically.

Inferences on dinosaur cognition are hindered by the fact that both extant crocodilians and birds are highly derived in their own ways: Convergently to mammals, birds have not only evolved an enlarged forebrain and cerebellum, but also extensive connections between these two brain regions (Gutierrez-Ibanez et al., 2018) and descending projections from the pallium to the brainstem and/or spinal cord (Ulinski and & Margoliash, 1990; Reiner & Medina, 2000). These circuits are likely essential for enabling various avian behaviors but are not present in extant non avian sauropsids (Ulinski and & Margoliash 1990; Gutierrez-Ibanez et al., 2023). It remains obscure when they evolved. Crown-group birds also possess an apomorphic dorsal projection of the telencephalon, the eminentia sagittalis or wulst, which appears to be absent even from endocasts of derived non-avian maniraptoriforms such as *Archaeopteryx* and *Stenonychosaurus* and is prominently involved in visual cognition (Walsh & Milner, 2011; Iwaniuk & Wylie, 2020). Crocodilians on the other hand conserve a plesiomorphic brain morphology and cerebral tissue organization (Briscoe et al., 2018; Briscoe & Ragsdale, 2018). They are unusual in being secondary ectotherms (e.g., Seymour et al., 2004; Legendre et al., 2016; Botha et al., 2023) and it is unclear how this might have affected their neurology and cognition. Thus, the extant archosaurian groups leave us in a rather suboptimal position to infer cognitive traits in non-avian dinosaurs.

Obviously, even the absence of a given cognitive trait in both crocodilians and basal extant birds like palaeognaths does not refute its existence in Mesozoic dinosaurs, considering the diversity and long evolutionary history of this group. In fact, a species’ ecology is typically more indicative of certain behaviors and associated cognitive phenomena than its phylogenetic affinities. For instance, habitual tool use is an adaptation typically found in omnivorous extractive foragers (Parker & Gibson, 1977; Parker, 2015) and is only rarely reported in predators (Shumaker et al., 2011). This is reflected by the fact that the most common types of tooling actions that have evolved comprise reaching, probing or pounding, usually in order to access food (Colbourne et al., 2021). It has long been observed that tool use appears when a species is found in an uncharacteristic niche, for which it lacks the appropriate morphological adaptations, and thus compensates by using tools to generate a functionally equivalent behavior (Alcock, 1972; Parker & Gibson, 1977). This is likely why a number of birds that use tools are found on islands, yet the ability appears absent in their close mainland relatives (Rutz et al., 2016). Simply put, in order for tool use to evolve, there needs to be a reason for it to evolve, and there are very few ecological contexts where tool use is a superior adaptation to its morphological equivalent (Hansell & Ruxton, 2008). Unfortunately, this type of extremely specific contextual information is nearly absent in long extinct species. From its iconic tooth and jaw morphology, one can confidently predict that a hypercarnivorous species like *T. rex* would have no need for tools, but the problem remains that few assumptions about extinct animal cognition are falsifiable.

In sum, reconstructing cognition in dinosaurs and other fossil taxa without close living analogs is a challenging endeavor that requires integrative approaches if we are to provide compelling inferences (de Sousa et al., 2023). Bare neuronal count estimates might be considered a rather minor contribution to this effort and need to be aligned with data from comparative anatomy and neurology, ecology, trace fossils, and comparative behavioral studies on extant animals to offer a plausible picture of cognition in extinct lineages. While communicating such findings, researchers should acknowledge the limitations of the presented inferences to allow their audience to delineate between reasoned conclusions and speculation. In a field such as dinosaur research - avidly followed by popular media and the public eye - a nuanced view appears especially warranted.

### Conclusions

The dinosaurian neuronal count and relative brain size estimates presented by Herculano-Houzel (2023) are inaccurate due to methodological shortcomings, in particular for *T. rex*. Accordingly, the biological inferences drawn from them are implausible. As we show here, there is no compelling evidence that relative brain size in large-bodied theropods differed significantly from that of extant non-avian sauropsids, and their telencephalic neuron counts were likely not exceptional, especially for animals of their size. Furthermore, we highlight issues associated with neuron count estimates in vertebrate paleontology and argue against their use in reconstructing behavioral and life history variables, especially in animals such as non-avian dinosaurs, for which disparate neuron densities might be hypothesized based on different phylogenetic and morphological arguments.

For obvious reasons, many inferences we might make about Mesozoic dinosaur behavior will remain limited. Nevertheless, we can justify certain predictions - to a degree - within integrative empirical frameworks to which neuron count estimates might well be added in the future. Before such steps can be taken, however, a substantially improved understanding of the relationship between neuron counts and other biological variables, especially cognitive performance, in extant animals is required.

## Institutional abbreviations

AMNH: American Museum of Natural History, New York City, New York, United States
BMNH / NHMUK: Natural History Museum, London, UK
BSP: Bayerische Staatssammlung für Paläontologie und historische Geologie, Munich, Germany
BYU: Brigham Young University, Earth Science Museum, Provo, Utah, United States
CAPPA/UFSM: Centro de Apoio à Pesquisa Paleontológica da Quarta Colônia / Universidade Federal de Santa Maria, São João do Polêsine, Rio Grande do Sul, Brazil.
CM: Carnegie Museum of Natural History, Pittsburgh, Pennsylvania
CMN: Canadian Museum of Nature, Ottawa, Ontario, Canada
DINO: Dinosaur National Monument, Jensen, Utah, United States
FIP: Florida Institute of Paleontology, Palm Beach, Florida, United States
FMNH: Field Museum of Natural History, Chicago, Illinois, United States
FPDM: Fukui Prefectural Dinosaur Museum, Fukui, Japan
HMN / MB.R.: Museum für Naturkunde, Berlin, Germany
IGM: Mongolian Institute of Geology, Ulaan Bator, Mongolia
IRSNB / RBINS: Institut Royal des Sciences Naturelles de Belgique, Brussels, Belgium
IVPP: Institute of Vertebrate Paleontology and Paleoanthropology, Beijing, China
KUVP: Kansas University Natural History Museum, Lawrence, Kansas, United States
MACN: Museo Argentino de Ciencias Naturales “Bernardino Rivadavia”, Buenos Aires, Argentina
MPC-D: Institute of Paleontology and Geology, Mongolian Academy of Sciences, Ulaan Bator, Mongolia
MUCPv-CH: Museo de la Universidad Nacional del Comahue, colección del Museo Ernesto Bachmann, Villa El Chocón, Argentina
MOR: Museum of the Rockies, Bozeman, Montana, United States
NMC: Canadian Museum of Nature, Ottawa, Canada
NCSM: North Carolina Museum of Natural Sciences, Raleigh, North Carolina, United States
OMNH: Sam Noble Museum at the University of Oklahoma, Norman, Oklahoma, United States
PIN: Paleontological Institute, Russian Academy of Sciences, Moscow, Russia
PKUP: Peking University Paleontological Collections, Beijing, China
ROM: Royal Ontario Museum, Toronto, Ontario, Canada
RTMP/TMP: Royal Tyrrell Museum of Palaeontology, Drumheller, Alberta, Canada
SGM: Ministere de l’Energie et des Mines, Rabat, Morocco
USNM: Smithsonian National Museum of Natural History, Washington, D.C., United States
UUVP: University of Utah, Salt Lake City, Utah, United States
YPM: Yale Peabody Museum, New Haven Connecticut, United States

## Acknowledgements

We want to thank David Burnham, Gregory M. Erickson, Ariana Paulina-Carabajal, and Lawrence M. Witmer for sharing valuable information on fossil specimens and Nicolas E. Campione for recommendations on body mass calculations. Andrew N. Iwaniuk is acknowledged for helpful discussions on the methodology and structure of the study and Jonathan Stone for allowing GRH to perform alligator dissections in his lab. Finally, we thank Stig Walsh and two anonymous reviewers for their thoughtful and constructive comments on earlier drafts of this manuscript.

## Supplementary Files

**Suppl. File 1**: Information on the selection and determination of brain endocast measurements in this study, including comments on endocast data presented in Jerison (1973), Hopson (1979) and Hurlburt (1996) and discussions on endocranial fill in non-maniraptoriform dinosaurs.

**Suppl. File 2**: Brain and body mass data of extant sauropsid species, excluding the olfactory tracts and bulbs. Brain mass data are derived from Hurlburt (1996), except for those of the Siamese crocodile, which derive from Chentanez et al. (1983). All birds considered here belong to non-telluravian taxa.

**Suppl. File 3**: Neuron count data, brain size measures and respective references for extant sauropsid and mammalian species.

