## Supplementary File 1 for "How smart was *T. rex*? Testing claims of exceptional cognition in dinosaurs and the application of neuron count estimates in palaeontological research"

**Part A: Discussion and amendments of endocranial volumes provided by Jerison (1973), Hopson (1979) and Hurlburt (1996).**

For our analyses, we (as well as Herculano-Houzel, 2023) relied on previously published endocranial volumes (EV) from three frequently referenced sources in the field of palaeoneurology: Jerison (1973), Hopson (1979) and Hurlburt (1996). All these authors used Graphic Double Integration (GDI) to arrive at their brain size estimates and only considered the "brain" endocast volume (BrEV) of the total endocast (excluding olfactory tract and lobes, the brainstem caudal to cranial nerve XII (Nervus hypoglossus), nerve or vessel stumps, the pituitary fossa, and/or the bony labyrinths; this procedure is followed here as well, see Materials and Methods). During the preparation of this work, we became aware of a number of inaccuracies or cases of intransparent reporting in these sources and took the opportunity to amend several EV estimates provided therein. Below, we detail our decisions to retain, re-calculate or discard EV values communicated by the aforementioned references, starting with Jerison (1973).

Jerison (1973) listed EV data for ten adult specimens of non-maniraptoriform dinosaur genera from the primary literature. We examined the original EV sources to identify specific epithets and identify specimens, neither of which were reported by Jerison (1973). For two genera, original EV data were reported in the cited sources (*Giraffatitan brancai* and *Stegosaurus ungulatus*); the other eight were calculated by Jerison (1973) using GDI. Jerison (1973) lists an EV of 310 mL for *Giraffatitan brancai* ("*Brachiosaurus*"), which was originally reported by Janensch (1935-36) who created a plasticine cast of the endocranial volume of a specimen now catalogued as MB.R.2223.1. This volume may have included parts of the olfactory tract and brain stem caudal to nerve XII and thus be a slight overestimate. We nevertheless retain this value here. For *Stegosaurus ungulatus*, Jerison (1973) used an EV (56 mL) reported by Gilmore (1914, not 1920) for an endocast of specimen YPM 1853. This artificial endocast was prepared by Marsh (1896). However, it was incomplete and Marsh did not indicate what parts had been restored (Galton, 2001). We discarded this value and instead used a GDI-derived EV from views of an undistorted and complete endocast of specimen CM 106 (Galton, 2001) which was previously reported by Hurlburt et al. (2013).

For the remaining eight genera, Jerison (1973) calculated the EV of the "brain portion" of the endocast (Figure 4) using GDI and assuming a brain mass:endocast volume ratio of 50%. He used dorsal and lateral views of photographs or drawings of endocasts of only five genera, of which we use two here, *Edmontosaurus annectens* ("*Anatosaurus*" in Jerison, 1973) YPM 618 (Lull and Wright, 1942), EV= 300 mL; and *Protoceratops andrewsi* AMNH 6466 (Brown & Schlaikjer,1940), EV = 30 mL. Note that the brain mass for *Protoceratops* was mistakenly reported as 28 g rather than 15 g by Hurlburt (1996), which would correspond to an intended 50% fill of the endocast volume. The erroneously inflated value was adopted by Herculano-Houzel (2023) who specifically remarked that the relative brain size of *Protoceratops* "approached the distribution of modern pre-K-Pg birds".

We did not consider Jerison's (1973) values for either *Iguanodon* NHMUK R2501 (Andrews, 1897), EV = 250 mL, *Triceratops* sp. (Hay, 1909), EV = 140 mL or *Tyrannosaurus rex* AMNH 5029 (Osborn, 1912), EV = 404 mL. We used RBINS R51 to represent *Iguanodon bernissartensis*. For this specimen, both stylopodial circumference measurements and CT-derived endocast data are available. For *T. rex* AMNH 5029, we relied on CT scans indicating an EV of 381.8 mL (Hurlburt et al., 2013). We also relied on CT-derived EVs for *Triceratops* sp., which endocranial morphology has received increased attention in recent years (Morhardt, 2016; Sakagami & Kawabe, 2020). We use the brain endocast volume of specimen MOR 1194, which was kindly shared by L. M. Witmer. The respective volume of ca. 230 mL (which corresponds reasonably well to EV data on specimen FPDM-V-9677 studied by Sakagami & Kawabe, 2020) is considerably larger than the 140 mL calculated by Jerison (1973) for specimen USNM 2416 (which was based on a plaster cast drawing in Hay, 1909). Jerison's (1973) still frequently cited estimate (see e.g., Button & Zanno, 2023) must thus be considered erroneous and should be ignored.

Two supposed GDI's from Jerison (1973) were based on lateral views only of the respective endocasts of *Allosaurus fragilis* AMNH 5753, EV = 335 mL; and *Diplodocus longus* AMNH 694 (a subadult individual - Witmer et al., 2008), EV = 100 mL (both from Osborn, 1912), which is inconsistent with the requirement of two orthogonal views to perform GDI. We relied on different values for these species. For *A. fragilis*, an EV of 98.5 mL was derived from GDI. Respective photographs were taken of an endocast replica of the specimen UUVP 294 (G. R. Hurlburt, private collection). For *Diplodocus*, information on the brain endocast volume of specimen CM 11161 was again provided by L. M. Witmer.

The final genus included by Jerison (1973) was *Camptosaurus* (EV 46 mL). We reject this value since there is no evidence of the existence of an actual endocast corresponding to the one possible source (Marsh, 1896: p. 197, Plate LXIII, Fig. 2). Jerison (1973) purportedly used GDI to find an EV of 46 mL for *Camptosaurus*. He cites Gilmore (1909), who figures no endocast but described *Camptosaurus* specimens discussed by Marsh (1896). The only possible GDI subject is a dorsal view of an endocast on a drawn reconstruction of the skull of "*Camptosaurus medius*" (= *C. dispar*) in Marsh (1896). Marsh's (1896) drawn skull reconstruction was based on isolated elements of the skull of "*Camptosaurus medius*" YPM 1880 and elements of the much larger skull of "*Camptosaurus amplus*" YPM 1887, which is now referred to *Theiophytalia kerri* (Brill & Carpenter, 2006). No sufficiently complete *Camptosaurus* skull was known in 1909, let alone 1896, from which an endocast could be derived from (Gilmore, 1909). Jerison's EV estimate for *Camptosaurus* must be discarded.

Hopson (1979) made original GDI-derived determinations of endocast volumes for *Kentrosaurus* (48 mL) and *Euoplocephalus* (82 mL) but provided no information about the specimens used. For reasons of transparency, we provide new GDI estimates here from illustrations of *Kentrosaurus aethiopicus* HMN Ki 124 (52.6 mL; figured by Galton, 1988) and *Euoplocephalus tutus* AMNH 5337 (82.7 mL; figured by Hopson, 1979). Interestingly, Hopson's estimates are closely approximating ours, suggesting Hopson's GDI was based on the same specimens.

Finally, we want to comment on EV values communicated by Hurlburt (1996). This reference provided an original brain volume estimate of 87.85 mL for *Ornithomimus edmontonicus* (= "*Dromiceiomimus brevitertius*"). However, this value was a GDI estimate derived from schematic depictions of an endocast included in Russell (1972). When based on photographs of the actual endocast (NMC 12228), GDI yields a significantly lower value of 49.89 mL which is applied here. Hurlburt (1996) also discussed brain size measurements for *Protoavis texensis*, which were originally provided by Chatterjee (1991) and later adopted by Herculano-Houzel (2023). Chatterjee (1991) reconstructed an artificial endocast out of flattened braincase elements from two specimens and derived a brain mass of 3.3 g from it. Hurlburt (1996) already questioned the legitimacy of Chatterjee's approach, suspecting that "there was an error in reconstructing the skull […] to produce what is probably a gross overestimate". In addition to that, the phylogenetic affinities of braincase material from *Protoavis* remain unresolved (Nesbitt et al., 2007), which further disqualifies its inclusion in comparative palaeoneurological studies.

**Part B: Details on the determination of EV in each specimen considered for analysis.**

We obtained EV data either from manual GDI performed on photographs or appropriate figures of endocasts from published descriptions or adopted reported endocast volumes from the literature. GDI was done by G.R.H., further details on the procedures are available upon request.

Where possible we used reported values for the "brain" region of endocasts (BrEV; see Materials and Methods). Where reported measures included the olfactory tract and lobes, the brainstem caudal to cranial nerve XII, nerve or vessel stumps, the pituitary fossa, and/or the bony labyrinth, we cautioned to exclude the volumes of these structures. In cases wherein authors had expressed the volumes of respective structures as percentages of the entire cast, we subtracted the same percentage. If this information was not provided, we used GDI on figures of the endocast to either determine the volume of the "brain portion" or else to determine the volume of the excluded region(s), which volumes were then subtracted accordingly.

**Maniraptoriformes**

For this group, we assumed that the brain filled 100% of BrEV (see Materials & Methods).

***Archaeopteryx lithographica*: **The endocast of BMNH 37001 with a total EV of 1.6 mL was used. EV was determined by CT. BrEV was 1.52 mL following subtraction of olfactory bulbs volume (0.077 mL). Data on total endocast volume was derived from Alonso et al. (2004) and on those of the olfactory bulbs from Balanoff et al. (2013).

***Bambiraptor feinbergi***: We used the EV of KUVP 129737, the endocast of the holotype of *Bambiraptor feinbergi* (AMNH FARB 30556), which represents a subadult animal presumably exhibiting adult brain size (see Materials & Methods). Data on BrEV derive from Hurlburt et al. (2013) and were originally determined by water displacement.

***Citipati osmolskae*****:** The endocast of IGM 100/978 with a total EV of 22.62 mL was used. EV was determined by CT. BrEV was 22.05 mL following subtraction of olfactory bulb volume (0.569 mL). Data from Balanoff et al. (2013).

***Khaan mckennai*****:** The endocast of IGM 100/973 with a total EV of 8.83 mL was used. EV was determined by CT. BrEV was 8.8 mL following subtraction of olfactory bulb volume (0.028 mL). Data from Balanoff et al. (2013).

***Ornithomimus edmontonicus*****:** We used GDI to calculate a total EV of 49.687 mL for a latex endocast mold of specimen NMC 12228, which lacks the olfactory bulbs as well as the pituitary fossa but also the ventralmost regions of the brain (Fig. S1). The calculated EV is thus an underestimate of the total brain volume.

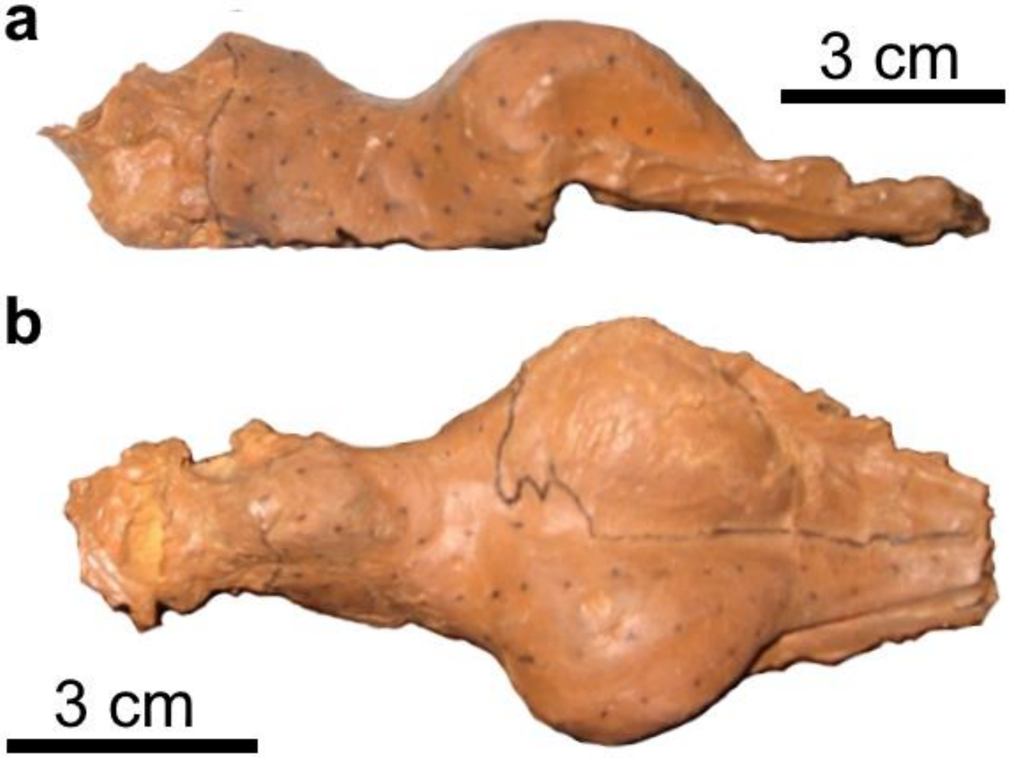


**Fig. S1**: Photographs of the latex endocast mold of specimen NMC 12228, assigned to *Ornithomimus edmontonicus*. **a**: Right lateral aspect. **b**: dorsal aspect.

***Shuvuuia deserti*****:** The endocast of IGM 100/977 with a total EV of 1.57 mL was used. EV was determined by CT. The EV for this specimen was initially reported to be 0.83 mL by Balanoff et al. (2013). However, a resegmentation of the respective braincase allowed this value to be corrected (Balanoff et al., 2024). BrEV was 1.52 mL following subtraction of olfactory bulb volume (0.05 mL). Data from Balanoff et al. (2013, 2024).

***Stenonychosaurus inequalis*****:** Currie & Zhao (1993) calculated a total EV of 45 mL by water displacement for a composite endocast (RTMP 1986.0036.457 & RTMP 1979.008.001) derived from braincase elements of "*Troodon formosus*". This name is now considered a *nomen dubium*, which is why we use *Stenonychosaurus inequalis* here. We also assume that

*Stenonychosaurus inequalis* and “*Latenivenatrix mcmasterae”* (to which RTMP 1979.008.001 has been assigned - van de Reest & Currie, 2017) are synonymous (Cullen et al., 2021). According to Currie & Zhao (1993), the provided EV might be a slight underestimate, given that the endocast did not include the large floccular lobes of the cerebellum and was moderately deformed due to crushing of left-side braincase elements. lacks the rostral half of the pituitary fossa However, more recently Morhardt (2016) generated a digitally restored composite endocast from the same braincase elements and provides an EV of only 39.74. Without, olfactory structures, the EV of the endocast is 38.65 mL (Morhardt, 2016). This estimate might still include cranial nerve stumps and small portions of the pituitary fossa (the rostral half of which is missing from the composite specimen – Morhardt, 2016). Erring on the side of a slight potential overestimate, we accept this value as the BrEV of *S. inequalis* for our study.

**Non-maniraptoriform Theropoda**

For this group, we assumed that the brain filled only a fraction of BrEV (see Materials & Methods). Two BrEV values were calculated for each endocast, one assuming that the brain occupied 42% of the respective endocast portion, and one assuming that it filled 31%.

***Acrocanthosaurus atokensis***: The CT-derived endocast of specimen OMNH 10146 was considered, which has been described by Franzosa & Rowe (2005). Total EV was determined to be 190.8 mL and included the bony labyrinth and pituitary. To extract BrEv, we performed GDI on Figure 2 of the respective paper and determined it to be 122.74 mL.

***Allosaurus fragilis***: We used GDI to obtain a BrEV of 98.5 mL from photographs of dorsal and left lateral views of a duplicate endocast of UUVP 294 (private collection of G.R.H; the original has been discussed by Rogers, 1998). We excluded the dorsal portion of the endocast from a line 0.405 cm ventral to its dorsal limit, as this does not appear to represent endocranial tissue. This estimate corresponds reasonably well to the CT-derived total EV of 163 mL reported for the conspecific specimen DINO2560 (Lessner et al., 2023).

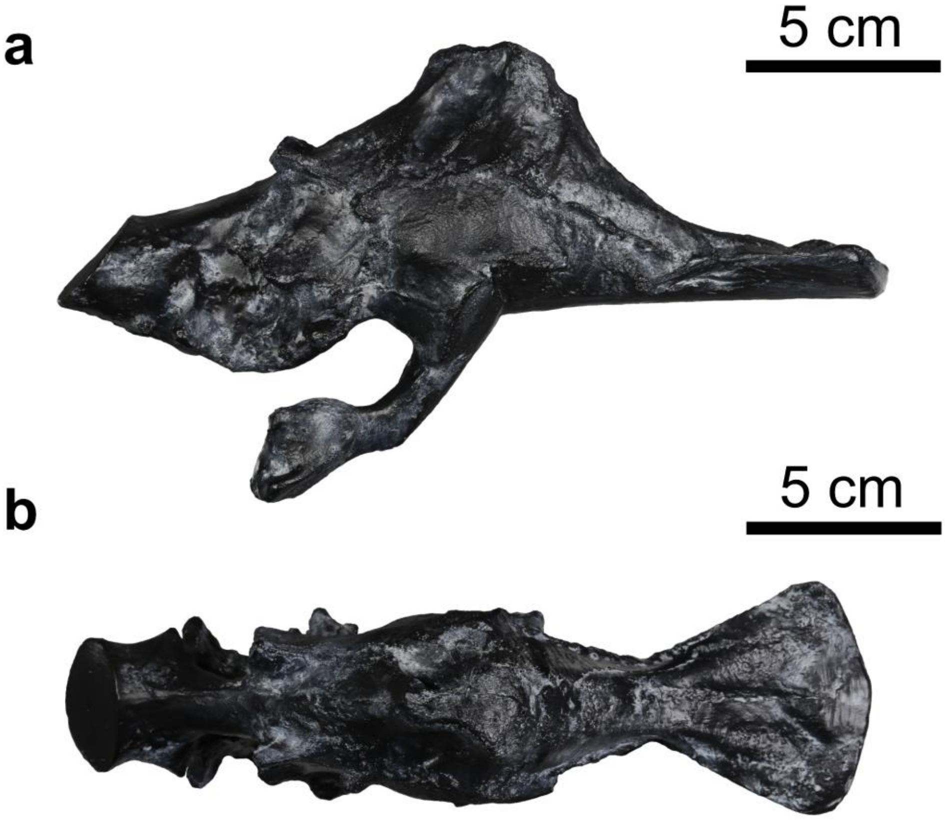


**Fig. S2**: Photographs of the duplicate of the natural endocast of specimen UUVP 294, assigned to *Allosaurus fragilis*. Note the prominent pituitary fossa. **a**: Right lateral aspect. **b**: dorsal aspect.

***Carcharodontosaurus saharicus*:** Larsson (2000) reported a BrEV (applying the same definition as we do herein) of 224.0 mL obtained by CT for specimen SGM-Din 1.

***Carnotaurus sastrei***: A CT-generated endocast for specimen MACN CH 894 with a total EV of 169.8 mL was described by Cerroni & Paulina-Carabajal (2019). We used GDI on Figure 2 of said paper to extract BrEV and determined it to be 108.34 mL. The scale bar in the respective figure measures 4 cm, not 5 cm as reported in the paper (verified by A. Paulina-Carabajal, pers. com.).

***Giganotosaurus carolinii***: Paulina- Carabajal & Canale (2010) determined a total EV of 275 mL for specimen MUCPv-Ch1 by water displacement (which equals a CT-derived EV communicated by Coria & Currie, 2003). This volume excludes nerve stumps, blood vessel casts resulting from foramen infills and the bony labyrinth. Applying the same definition used herein, the authors determined a BrEV of 225 mL, a value we adopted for our analyses.

***Majungasaurus crenatissimus***: A CT-generated endocast for specimen FMNH PR 2100 with a total EV of 117.16 mL was described by Sampson & Witmer (2007). Total EV included the olfactory tracts and bulbs as well as probably the pituitary volume, although this was not stated. We assume that this measure excluded the bony labyrinth. We used GDI on Figure 18 of said paper to extract BrEV and determined it to be 89.32 mL. Morhardt (2016) generated a digitally restored endocast from the same specimen and provides an EV of 106.09 mL, not considering the olfactory tract and bulbs but apparently including the pituitary and cranial nerve stumps.

***Sinraptor dongi***: Paulina-Carabajal & Currie (2012) described a CT-generated endocast from specimen TMP 93.115.1 (cast of IVPP 10600) and extracted a BrEV (defined as herein) of 95 mL. This measurement includes minor parts of the pituitary and cranial nerve stumps (A. Paulina-Carabajal, pers. com.) so that it represents a slight but tolerable overestimate. We assumed that the specimen represents an adult individual (see Materials & Methods).

***Tarbosaurus bataar***: We considered the artificial latex endocast of specimen PIN, no. 553-3/1 (Saveliev & Alifanov, 2007). This cast only captures the right side of the endocranial cavity. Total EV was estimated by Saveliev & Aifanov (2007) to be.184 mL. Since only a lateral view of the incomplete cast was provided, we used comparative anatomical data from its closest known relative, *Tyrannosaurus rex* (Brusatte & Carr, 2016), to extract BrEV from total EV. In *T. rex* AMNH 5029, BrEV equals 86.53% of total EV (when excluding the pituitary volume, which is not preserved in PIN, no. 553-3/1). Applying this ratio to the *Tarbosaurus* specimen, a BrEV of 159.2 mL.

***Tyrannosaurus rex***: BrEV measurements (following the same definition as applied herein) for all three specimens included here (FMNH PR 2081, AMNH 5029, AMNH FR 5117) were derived from CT data reported by Hurlburt et al. (2013).

**Non-theropod dinosaurs (Sauropodomorpha & Ornithischia)**

For this group, we assumed that the brain filled only a fraction of BrEV (see Materials & Methods). Two BrEV values were calculated for each endocast, one assuming that the brain occupied 42% of the respective endocast portion, and one assuming that it filled 31%.

***Amargasaurus cazaui:*** Paulina-Carabajal et al. (2014) describe the CT-derived endocast of specimen MACN-N 15, which we include here. *Amargasaurus* is a dicraeosaurid, a sauropod family which endocasts are characterized by a conspicuous dorsal dural expansion, which was not occupied by neural tissue (Janensch, 1935-36; Paulina-Carabajal et al., 2014). The total EV of the specimen including the dorsal sinus is 150 mL and 94-98 mL without it, while still including cranial nerve and vascular infillings (Paulina-Carabajal et al., 2014; A. Paulina-Carabajal, pers. com.). We applied GDI to extract BrEV, arriving at a volume of 84 mL.

***Apatosaurus* sp.*:*** The CT-derived endocast of BYU 17096, which was described by Balanoff et al. (2010) was considered. The species status of this specimen has not been clarified, for the respective body mass estimate, we rely on a specimen of *A. lousiae*. Our body mass estimate. The total EV was reported to be 125.14 mL, the olfactory bulbs are not preserved but this estimate includes the olfactory tracts, pituitary and extensive nerve trunks. We applied GDI to extract BrEV, arriving at a volume of 102.48 mL.

***Buriolestes schultzi***: A CT-derived endocast for specimen CAPPA/UFSM 0035 of this basal sauropodomorph was described by Müller et al. (2021). The total EV was given as 2.8 mL. We extracted BrEV using GDI and determined it to be 2.43 mL. The reported length for the scale bar in Fig. 8 D-E was indicated as 20 mm whereas it appears to have been approximately 25 mm, when compared to the linear measurements from the endocast reported in the paper. Therefore, the latter were used to for the determination of BrEV.

***Diplodocus* sp.*:*** We used the virtual endocast of CM 11161, which was generated via CT. The taxonomic placement of this skull is controversial. It is typically assigned to *Diplodocus longus*, a species based on highly fragmentary remains that do not include cranial material. We thus follow the recommendation of Tschopp et al. (2015) and refer to CM 11161 as *Diplodocus* sp. here. Our respective body mass estimate derives from an individual assigned to *Diplodocus carnegiei*. Total EV of the endocast is 106.4 mL (L. M. Witmer, pers. com.). We subtracted 6% of total EV to account for the pituitary fossa (which corresponds to 6% of total EV in *Amargasaurus*, another diplodocoid taxon - Paulina-Carabajal et al. 2014), leaving us with a BrEV of 100 mL. This could still be a slight overestimate, given that the incompletely preserved olfactory bulbs (Witmer et al., 2008) as well as nerve trunks may still be included in this measure.

***Edmontosaurus annectens***: Lull and Wright (1942) described the endocast of specimen YPM 618 (formerly "*Anatosaurus*" now referred to *Edmontosaurus annectens* - Xing et al., 2014) and determined a total EV of 450 mL by water displacement. We rely on the GDI-derived BrEV of 300 mL determined by Jerison (1973).

***Euoplocephalus tutus***: We relied on the illustration of the endocast of specimen AMNH 5337 from Hopson (1979) to derive BrEV by means of GDI, determining it to be 82.7 mL. This very closely matched the findings of Morhardt (2016), who determined the EV of the conspecific specimen AMNH FR 5405 to be 83.08 mL (excluding olfactory structures, total EV = 89.96).

***Giraffatitan brancai***: The artificial plasticine endocast by Janensch (1935-36) of the endocranial cavity of MB.R.2223.1 (formerly known as HMN t1 or simply T1) with a total EV of 310 mL was used. We adopt this as the BrEV of the specimen here. While this volume does not include the pituitary fossa and the bony labyrinth, other structures we exclude from BrEV are covered by this estimate, including the olfactory tracts. Based on the descriptions of Janensch (1935-36) we assume that care was taken to exclude nerve and blood vessel foramina in the measurement. Thus, the BrEV considered here must be considered an overestimate, but a tolerable one. This is especially true since the respective skull, though adult, does not represent the largest known one of this species (Janensch, 1935-36).

***Hypacrosaurus altispinus***: The endocast of ROM 1247 with a total EV of 289.9 mL was used. EV was determined by CT. BrEV was 275.9 mL following subtraction of olfactory bulb and tract volume (14 mL). We assumed that BrEV did not include nerve stumps and the bony labyrinth although this was not stated. Hence, the BrEV considered here might be an overestimate. Data from Evans et al. (2009).

***Iguanodon bernissartensis***: The CT-derived endocast of RBINS R51, which was described by Lauters et al. (2012) was included. The authors note that the total EV excluding the olfactory system equals 357 mL. We adopt this value as BrEV here. The lateral view of the endocast which is presented in Fig. 16.3 of the respective paper does not depict well demarcated blood vessels, cranial nerves, the pituitary gland, or semicircular canals. However, the endocast does include portions of the spinal cord caudal to the hypoglossal nerve, so that BrEV certainly is somewhat exaggerated. Since no appropriate depictions of the endocast were presented, DGI could not be applied.

***Kentrosaurus aethiopicus***: We relied on the illustration of the endocast of specimen HMN Ki 124 from Galton (1988) to derive BrEV by means of GDI, determining it to be 52.6 mL.

***Protoceratops andrewsi***: Brown & Schlaikjer (1940) described and figured the plaster endocast of specimen AMNH 6466. We rely on the GDI-derived BrEV of 30 mL determined by Jerison (1973).

***Psittacosaurus lujiatunensis*:** The endocranial anatomy of *P. lujiatunensis* was studied by Zhou et al. (2007). We consider the BrEV (defined by Zhou et al. 2007 as done here) provided for the largest of the studied specimens, PKUP V1060, which is 14.3 mL. We are not aware of published femoral circumference data for adult *P. lujiatunensis*. For our body mass estimate, we relied on specimen AMNH 6541, which is an adult *Psittacosaurus* of undetermined species status. The length of this specimen's femur (178.7 mm) falls well in the range of those of adult *P. lujiatunensis* (Zhao et al., 2013). The plausible body mass range calculated for AMNH 6541 by us (21.3 – 35.9 kg) covers the 25 kg estimate for PKUP V1060 that is cited by Zhou et al. (2007) and which was "estimated from femoral dimensions" by aid of an undisclosed method.

***Stegosaurus ungulatus***: The BrEV measurement of 64.2 mL (following the same definition as applied herein) for specimen CM 106 (figured in Galton, 2001) was adopted from Hurlburt et al. (2013), who determined it via GDI.

***Thescelosaurus neglectus***: The CT-derived endocast of NCSM 15728 with a total EV of 30.43 mL was used. The pallial portion of the endocast was distorted und digitally restored. This resulted in slightly divergent minimum and maximum estimates for the specimen's total EV (29.07 mL compared to 30.43 mL). Erring on the side of a potential overestimate, we considered the larger one of these values. Hence, BrEV was 27.65 mL following subtraction of olfactory bulb and tract volume (2.78 mL). Data from Button & Zanno (2023).

***Triceratops* sp.**: We relied on a CT-derived BrEV for MOR 1194 provided by L. M. Witmer, which excludes volumes of the pituitary fossa, olfactory structures, the bony labyrinth and cranial nerve stumps. The latter are notably massive in *Triceratops* (see Sakagami & Kawabe, 2020). The specimen is an isolated adult braincase that has been previously studied and discussed by Morhardt (2016). BrEV is 228.75 mL. As previously mentioned, the BrEV of 140 mL provided by Jerison (1973) based on an illustration of the plaster endocast of specimen USNM 2416 in Hay (1909) must be considered erroneous.

**Part C: Additional comments on brain:endocast volume ratios in selected non-maniraptoriform dinosaurs.**

We justified a brain:endocast volume ratios of 31-42% in dinosaurs based on comparative evidence from modern crocodilians. We explicitly exclude the taxa Maniraptoriformes and Pachycephalosauria here, for which extensive anatomical evidence suggests increased endocranial fills (see main text and references therein). Nevertheless, it has been argued, that some dinosaurian groups outside of these groups exhibited endocranial fills significantly exceeding 50%, especially the hadrosauriform ornithischians (Evans, 2005; Evans et al., 2009; Knoll et al., 2021), of which we consider three genera (*Edmontosaurus*, *Iguanodon*, *Hypacrosaurus*) in our analyses. However, we are not compelled by the arguments brought forward so far to justify this assumption.

The primary anatomical evidence for a secondarily increased endocranial fill in these dinosaurs is the presence of branching vascular imprints, so-called valleculae, on the endocranial surface adjacent to the lateral poles of the cerebrum (Evans, 2005; Lauters et al., 2013), potentially suggesting a comparatively tight fit of the brain and its associated blood vessels in the braincase. Interestingly, similar structures have also been identified in tyrannosaurid theropods: The braincases of *Tarbosaurus* and *Tyrannosaurus* show networks of valleculae in the same anatomical region (Witmer & Ridgely, 2009). These patterns have been taken to correspond to blood vessels pressed against the endocranial surface by the brain through the meninges (Witmer & Ridgely, 2009). Such an interpretation is supported by observations on modern American alligators (*Alligator mississippiensis*). In these animals, ramifying meningeal arteries occur on the dura external to the lateral cerebrum where it bulges against the arachnoid and dura mater (Fig. 2B).

There is a second patch of valleculae on the endocranium adjacent to a region of the lateral brain caudal to the cerebrum in *T. rex* (Witmer & Ridgely, 2009). A patch of ridges in the same location, caudal to the cerebrum, has also been reported for a hadrosaur endocast (Evans, 2005). In this region of the alligator brain, a layer of cerebrospinal fluid lies between the arachnoid and the midbrain, which does not press laterally onto the meninges or endocranium (Fig. 2). Accordingly, these "midbrain" valleculae would have corresponded to (probably arterial) meningeal vessels, indicating that the previously mentioned valleculae lateral to the cerebrum were also associated with superficial meningeal vessels and not vessels running directly along the brain's surface. Thus, vascular imprints in the aforementioned groups do not necessarily indicate endocranial fills that exceed those of other dinosaurs or extant crocodilians.
